# Rapid covalent labeling of a GPCR on living cells using a nanobody-epitope tag pair to interrogate receptor pharmacology

**DOI:** 10.1101/2022.05.09.491166

**Authors:** Chino C. Cabalteja, Ross W. Cheloha

## Abstract

Synthetic molecules that form a covalent bond upon binding to a targeted biomolecule (proximity-induced reactivity) are the subject of intense biomedical interest for the unique pharmacological properties imparted by irreversible binding. However, off-target covalent labeling and the lack of molecules with sufficient specificity limit more widespread applications. We describe the first example of a crosslinking platform that uses a synthetic peptide epitope and a single domain antibody (or nanobody) pair to form a covalent linkage rapidly and specifically. The rate of the crosslinking reaction between peptide and nanobody is faster than most other biocompatible crosslinking reactions, and it can be used to label live cells expressing receptor-nanobody fusions. The rapid kinetics of this system allowed us to probe the consequences on signaling for ligand crosslinking to the A2A-adenosine receptor. Our method may be generally useful to site-specifically link synthetic molecules to receptors on mammalian cell surfaces.

## Introduction

Molecules that form a covalent bond upon binding to their biological target have been applied as pharmacologically active agents for decades^1^. The reactivity of these molecules depends upon binding-mediated juxtaposition of reactive groups, a phenomenon referred to as proximity-induced reactivity (PIR). In contrast to molecules that bind to target proteins via transient non-covalent interactions, PIR molecules form a covalent crosslink after binding that usually persists throughout the biological lifetime of the target. These permanent modifications often lead to profound and enduring physiological responses when applied *in vivo*^2^. However, off-target reactions between PIRs and other proteins can have negative and irreversible biological consequences^3^. It is therefore critical for PIR molecules to interact with a very high level of specificity with their target.

Relative to small molecules, which are often used as the scaffold for the design of PIR compounds, peptides and proteins can be designed with better specificity for a protein target^4^. The introduction of crosslinking moieties into protein-specific polypeptide binders offers an appealing path to highly specific PIR compounds^5^; however, there are challenges with this approach. Any crosslinking moiety that is used must not interfere with target binding or induce off-target binding. Next, the crosslinking moiety must react with amino acids found at the binding site within the target protein and avoid reacting with amino acids in the peptide portion of the PIR compound itself. PIR compounds are often used to target Cys residues as they are relatively rare and show unique reactivity^6–11^, but this approach is limited when targeting proteins on cell surfaces as the Cys residues are typically engaged in disulfide bonds and therefore not available for crosslinking. Crosslinking moieties that react with other amino acids such as Lys, His, and Tyr have been used to crosslink peptide-based PIR compounds,^12–20^ and innovative strategies to simultaneously optimize both polypeptide sequence and crosslinking properties have also been developed^11^.

Among polypeptide scaffolds used to specifically label target proteins of interest, antibodies (Abs) are the most widely used. Methods to covalently link antibodies to their target through PIR would be valuable, but there are currently only a few examples. Antibodies that bind to small molecule haptens have been modified using hapten-crosslinker conjugates^21,22^. In a related example, a viral peptide epitope was modified with sulfonyl (VI) fluoride exchange (SufEx) electrophiles to facilitate crosslinking with antibodies that bound to that peptide^23^. Single domain antibody fragments (nanobodies, Nbs)^24^ have been converted to PIR compounds through the incorporation of non-natural crosslinking amino acids via translational reprogramming^13–15^. In each of these cases the Abs or Nbs that covalently bound to their target showed new or improved properties relative to the non-crosslinking versions. However, most of these strategies are not easily translated to other target proteins, and the crosslinking kinetics are relatively slow (k_2_ ~ 10^3^ M^-1^ s^-1^, labeling t_1/2_ > 1 h)^13–15^. Rapid labeling is especially important for methods to study the highly dynamic signaling responses initiated upon activation of cell surface proteins (t_1/2_ ~seconds to minutes). To address these issues, we have developed a peptide-based PIR compound derived from a 14-amino acid peptide (**6E** peptide)^25^ that is recognized by a Nb (Nb_6E_), previously shown to be amenable for use as an epitope tag^26^. Using this system, we demonstrate selective and rapid (k_2_ ~ 10^5^ M^-1^ s^-1^, >50-fold faster than previous reports) covalent labeling of nanobody and nanobody-receptor fusions expressed on live cells. We demonstrate that our new methodology can be used to dissect the impact of covalent tethering on agonist-induced GPCR signaling.

## Results

The crosslinking moiety was placed at either position 5 or 14 of the peptide based on a previous study of the structure-activity relationships for the binding of Nb_6E_ to analogues of the 6E peptide (Figure 1A)^27^. To avoid autoreactivity, we used variants of 6E in which the natural Lys at position 7 was replaced with Ala and the N-terminus was acetylated (Figure 1B, see Supporting Figure 1). Previous work showed that modifications at positions 7 and 11 cause minor changes in binding properties, whereas modification at position 5 causes a modest reduction in binding^27^.

**Figure 1.**
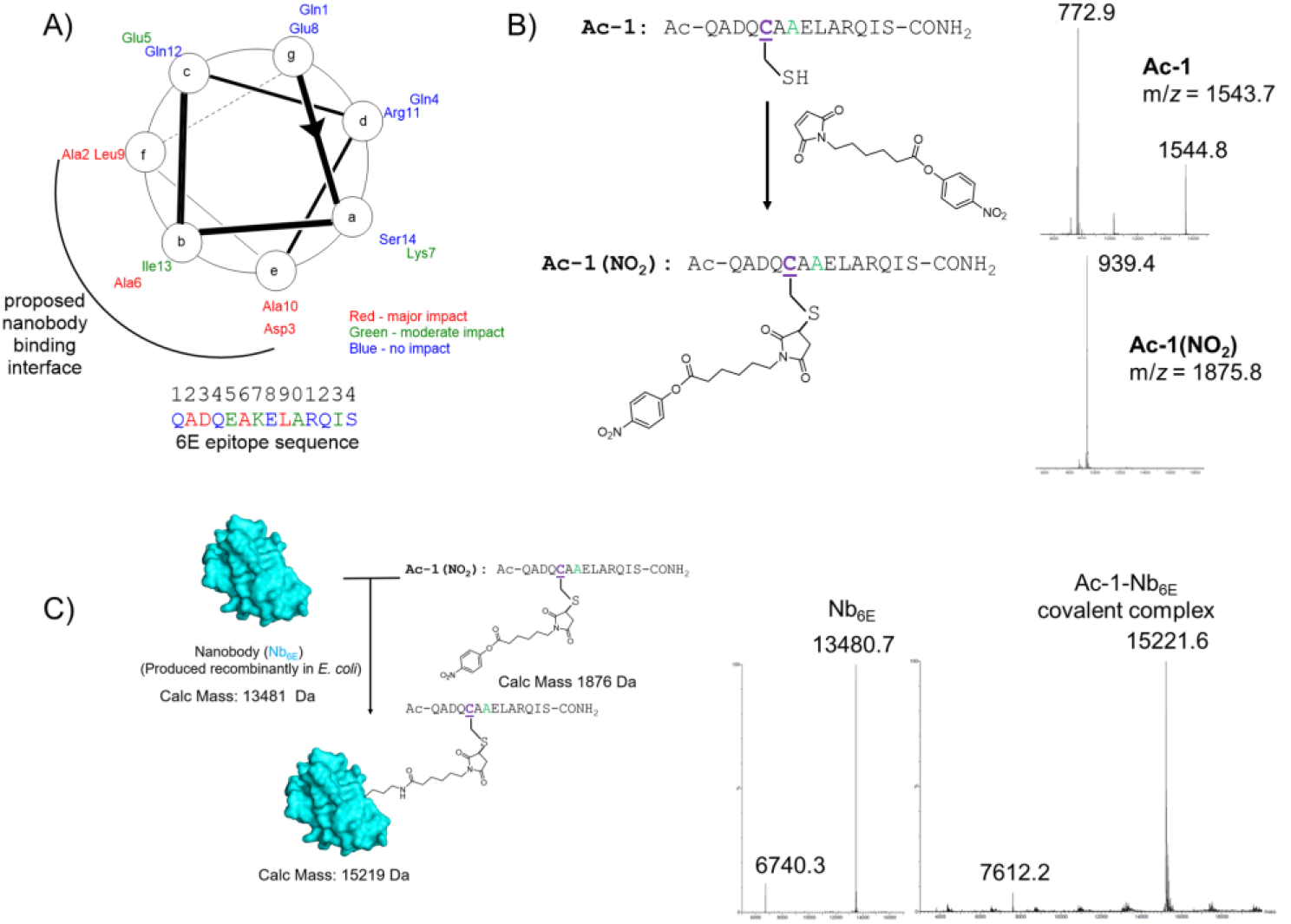
Design, synthesis, and preliminary evaluation of the Nb_6E_-6E epitope crosslinking pair. A) Summary of structure-activity relationship studies of the interaction between Nb_6E_ and the 6E peptide epitope^27^. Amino acid residues are mapped onto a hypothetical α-helical wheel diagram and a putative interface for binding to Nb_6E_ is illustrated. B) Synthetic scheme for preparation of crosslinking peptide-electrophile conjugates using cysteine-maleimide chemistry. Sites with mutations relative to the original peptide sequence are shown in colored font. Mass spectrometry data from LC/MS is shown at right. The full set of MS data is shown in Supporting Table 1. C) Schematic reaction between Nb_6E_ and **Ac-1(NO_2_)**. Deconvoluted MS data for Nb_6E_ (5 μM) before and after incubation with **Ac-1(NO_2_)** (20 μM) for 1 hour is shown at right.

For the crosslinking moiety, we chose activated phenol esters (see Figure 1 and Supporting Figure 1 for structures) which are known to crosslink with Lys and His sidechains^28,29^. We connected the crosslinking moieties to 6E peptides using cysteine-maleimide chemistry followed by purification with HPLC (Figure 1B). Preliminary kinetic crosslinking data (Supporting Figure 2A-D), acquired using SDS-PAGE and mass spectrometry, showed that every peptide-crosslinker conjugate reacted with Nb_6E_ to form a product with a higher molecular weight. One compound (**Ac-1(NO_2_)**) reacted the fastest with Nb_6E_ to form a singly modified product, a hallmark of PIR (Figure 1C). To test for specificity, **Ac-1(NO_2_)** was incubated with an unrelated Nb. This negative control reaction showed no crosslinking even after overnight incubation. (Supporting Figure 3). Analysis of the crosslinking reaction between Nb_6E_ and **Ac-1(NO_2_)** using trypsin digestion and mass spectrometry suggested that several different residues from Nb_6E_ might react to form a crosslinked product with **Ac-1(NO_2_)** (Supporting Figure 4). Attempts to structurally characterize the crosslinked 6E-Nb_6E_ complex (Ac-1-Nb_6E_) using crystallography were unsuccessful due to a lack of suitable crystals after screening >1500 conditions (data not shown). **Ac-1(NO_2_)** was stable in phosphate buffered saline in the absence of crosslinking Nb (half-life >20h, Supporting Figure 5).

Next, we characterized the kinetics of the reaction between crosslinking peptides and Nb_6E_. Upon mixing Nb_6E_ (5 μM) with an excess of crosslinking peptide (20 μM), conversion to product was monitored using either mass spectrometry or SDS-PAGE (Figure 2A, Supporting Figure 2). From these measurements, pseudo first-order rate constants were determined for the reaction that crosslinks the peptide and Nb_6E_ (Figure 2A). Affinity and reaction kinetics were also measured for the reaction between Nb_6E_ and fluorescein-labeled peptide (**FAM-1(NO_2_)**) using in-gel fluorescence (Figure 2B). Using this method, we calculated a K_D_ of 210 nM and a second order rate constant of 97,000 (M^-1^, sec^-1^). The rate of this reaction compares favorably with many other commonly used methods for successful biological conjugation, and the kinetics are comparable to enzymatic labeling using single domain enzyme tags such as SNAP and HALO tag^30,31^ (Figure 2C). Another estimate of peptide affinity was determined using surface plasmon resonance to measure the binding of a peptide without the crosslinking group (Ac-1(FAM), see Supporting Figure 1 for structure) to Nb_6E_. These experiments provided an estimate for **Ac-1(NO_2_)** binding (Supporting Figure 6) that is within the same order of magnitude as the data above (Figure 2B).

**Figure 2.**
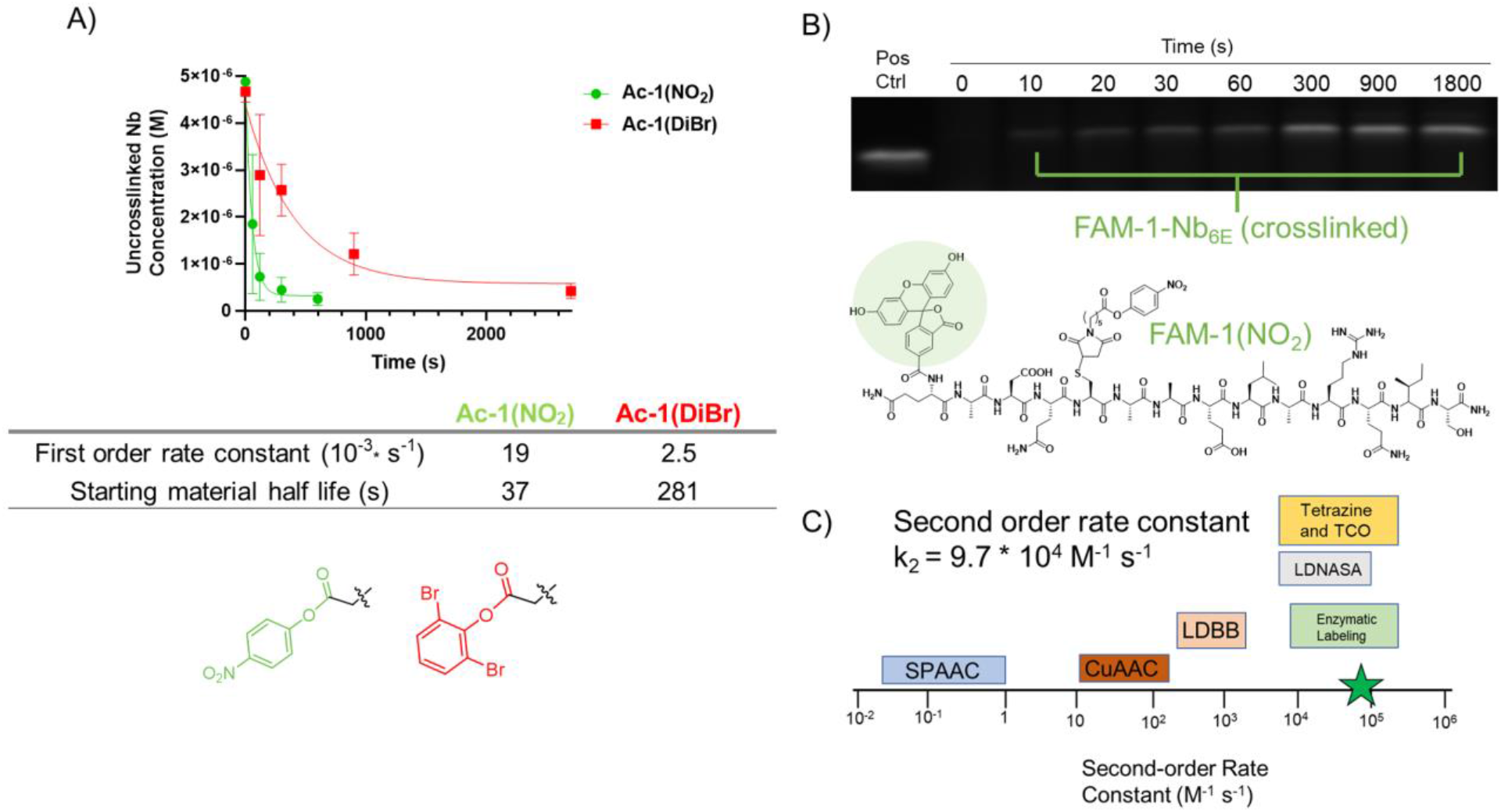
Evaluation of epitope-nanobody crosslinking kinetics. A) Measurement of the rate of reaction between Nb_6E_ and crosslinking peptides with varying ester groups (see supporting Figure 1 for full chemical structures). Nb_6E_ (5 μM) was mixed with an excess of the indicated crosslinking peptides (20 μM). The abundance of remaining non-crosslinked Nb_6E_ was quantified using LC/MS. B) Assessment of crosslinking kinetics using in-gel fluorescence. Nb_6E_ (100 nM) was mixed with an excess of **FAM-1(NO_2_)** (5 μM). Crosslinking was quenched at the indicated time points by addition of an excess of unlabeled competitor 6E peptide. Samples were run on SDS-PAGE and formation of the crosslinked product was assessed by in-gel fluorescence with quantitation by densitometry. Additional data from all in-gel fluorescence experiments is found in Supporting Figure 2E. C) Comparison of second-order rate constants for different types of chemoselective reactions. Rate constant ranges are adapted from previous work^30^. SPAAC: strain-promoted azide alkyne cycloaddition. CuAAC: copper-catalyzed azide alkyne cycloaddition reaction. LDBB: ligand-directed dibromophenyl benzoate reaction. Tetrazine and TCO: transcyclooctene and tetrazine reaction. LDNASA: ligand-directed N-acyl-N-alkyl sulfonamide reaction. Enzymatic labeling: SNAP tag and Halo tag reactions. The rate of the reaction characterized in panel B (97,000 M^-1-^s^-1^) is marked with the green star.

Our next goal was to apply the crosslinking reaction between **Ac-1(NO_2_)** and Nb_6E_ to label a receptor-nanobody fusion construct in which Nb_6E_ is localized to the extracellular side of the plasma membrane. We developed a HEK293 derived cell line stably expressing the A2A adenosine receptor, which is a member of the G-protein coupled receptor (GPCR) superfamily. This receptor was genetically fused at its N-terminus to Nb_6E_ (A2AR(Nb_6E_)), (Figure 3A) as well as an epitope tag (Alfa tag) added for detection with an orthogonal nanobody (Nb_Alfa_, see Supporting Methods for receptor sequence information)^32^. To assess binding, we used flow cytometry and 6E peptides functionalized at their N-terminus with fluorescein for detection (**FAM-1(NO_2_)** and **FAM-1**, see structures in Figure 2 and Supporting Figure 1). Incubation of A2AR(Nb_6E_)-HEK cells with either crosslinking or non-crosslinking peptide resulted in cellular staining consistent with binding to the receptor (Figure 3B, Supporting Figure 7). This staining was absent in control HEK293 cells that were not transfected with A2AR(Nb_6E_) (Figure 3B, green trace, Supporting Figure 7), which demonstrates high specificity for **FAM**-**1(NO_2_)** labeling of A2AR(Nb_6E_). To distinguish non-covalent binding of (expected mode of labeling for **FAM-1**) from covalent modification (expected for **FAM-1(NO_2_)**) an unlabeled peptide was used as a competitor to test whether the fluorescein-labeled peptides could be displaced from the receptor after binding (Figure 3B). As expected, the unlabeled peptide successfully displaced **FAM-1** from the cells, but it did not impact cells treated with **FAM-1(NO_2_)** (Figure 3B).

**Figure 3.**
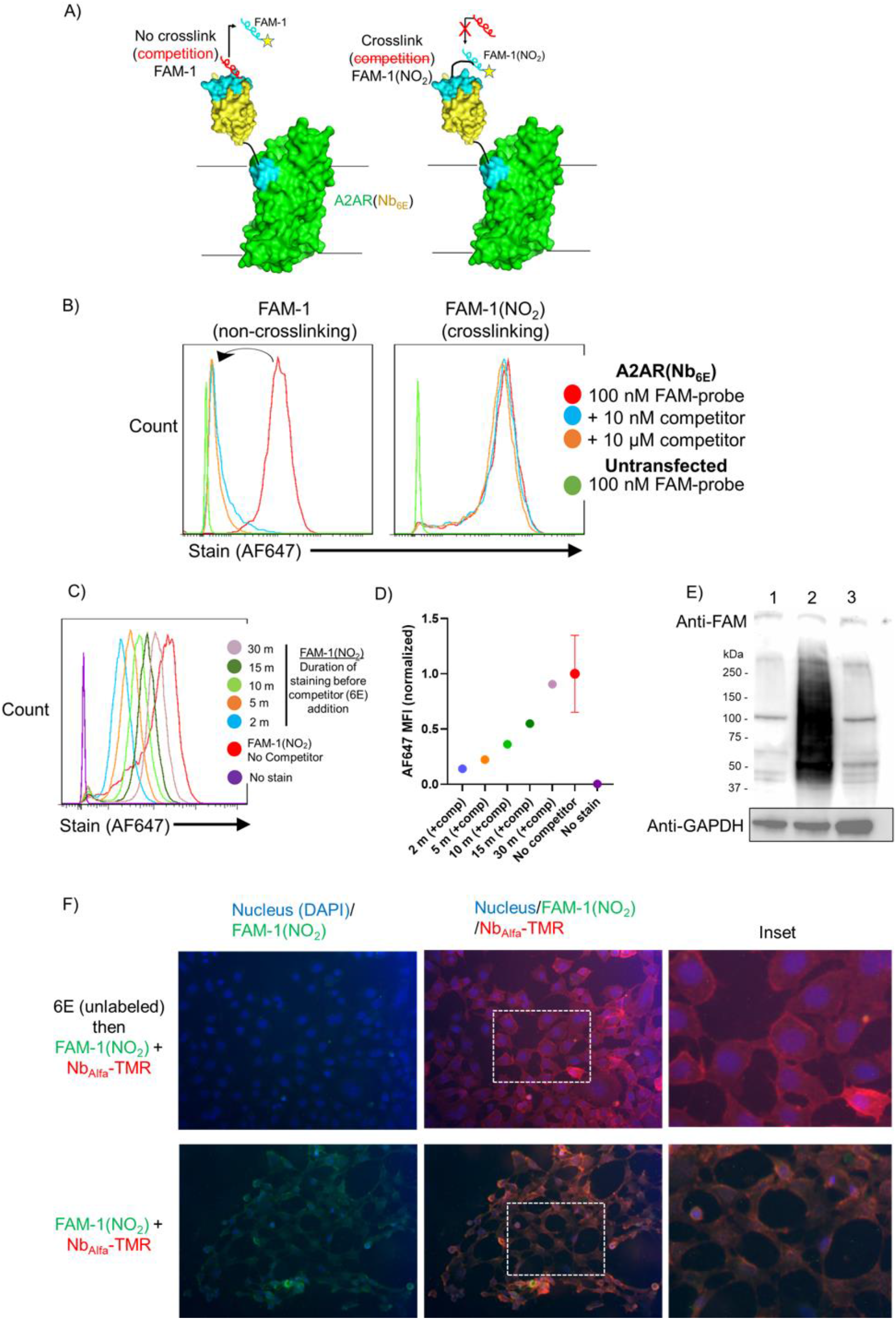
Evaluation of epitope-nanobody crosslinking on mammalian cells. A) Schematic of A2AR(Nb_6E_) binding to labeled 6E peptide (cyan) or unlabeled competitor peptide (red). The model of receptor is hypothetical and prepared using Pymol and a published crystal structure (PDB: 4UG2). See Supporting Methods for sequence information of this receptor. B) Comparison of cells expressing A2AR(Nb_6E_) treated with **FAM-1** (left) or **FAM-1(NO_2_)** (right). Cells were treated with 100 nM of fluorescein-labeled peptides on ice for 30 m, washed, and then exposed to unlabeled competitor peptide (6E) at the indicated concentrations for an additional 30 m. After washing, FAM-labeled peptides were detected with Alexafluor647 (AF647) modified anti-FAM antibody and analyzed by flow cytometry. C) Kinetics of covalent labeling of cells expressing A2AR(Nb_6E_). Cells were labeled with 100 nM of **FAM-1(NO_2_)** for the indicated durations at which time unlabeled competitor peptide was added to prevent further labeling. Total incubation time (before and after addition of competitor) is the same for all samples (30 m total). Cells were washed and analyzed as in panel B. “No stain” indicates no FAM-labeled peptide was added. “No competitor” indicates no unlabeled competitor peptide was added. D) Quantitation of data from panel C. Data correspond to mean ± SD from three independent experiments. E) Analysis of probe labeling specificity using Western Blotting. Lane 1: No probe added; Lane 2: 100 nM **FAM-1(NO_2_)**; Lane 3: Pretreatment with 10 μM unlabeled competitor peptide (6E) followed by treatment with 100 nM **FAM-1(NO_2_)**. Samples were resolved by SDS-PAGE and transferred to PDVF membranes. Labeled proteins were detected using horseradish peroxidase (HRP) conjugated anti-FAM antibody or HRP-anti-GAPDH. F) Analysis of crosslinking peptide labeling using fluorescence microscopy. HEK293 cells expressing A2AR(Nb_6E_) were stained with **FAM-1(NO_2_)** (100 nM) and Nb_Alfa_-TMR (300 nM) either with (top) or without (bottom) pre-treatment with unlabeled competitor 6E peptide. Cells were washed, fixed, stained with DAPI and imaged. Left and middle panels show identical images with filter sets applied as indicated. Right panels show the boxed inset from the middle panel.

To evaluate whether covalent labeling could be extended to a different receptor, we generated a cell line stably expressing a different GPCR (parathyroid hormone receptor-1, PTHR1)^33^ fused to Nb_6E_ within the N-terminus, also found at the extracellular face of the receptor (Nb_6E_-E2-PTHR1, Supporting Figure 8). Cells expressing Nb_6E_-E2-PTHR1 were labeled with **FAM-1(NO_2_)** and that labeling was not susceptible to displacement with unlabeled competitor peptide, in line with observations for cells expressing A2AR(Nb_6E_). In cell signaling assays, Nb_6E_-E2-PTHR1 was activated by the prototypical PTHR1 ligand PTH(1-34) with a potency comparable to past work using wild-type PTHR1 (Supporting Figure 9)^26^. These observations provide evidence that Nb_6E_-GPCR fusions can fold correctly, traffic to the cell surface, and respond to their natural ligands.

The kinetics of the covalent reaction between **FAM-1(NO_2_)** and A2AR(Nb_6E_)-HEK cells was measured by addition of the unlabeled competitor peptide at different times during the reaction to halt further crosslinking and to eliminate contributions from non-covalent binding (Figure 3C-D). The results from this experiment showed that the covalent crosslinking reaction proceed rapidly, with maximal levels of staining observed within 30 minutes after addition of **FAM-1(NO_2_)**. This reaction rate is consistent with the kinetics experiments performed in solution. To demonstrate binding specificity, we analyzed Western blots of intact cells exposed to **FAM-1(NO_2_)** that were washed and then lysed (Figure 3E, Supporting Figure 10). These data show a smeared, high molecular weight band present in cell lysates after treatment with **FAM-1(NO_2_)**, but the same band is not present in untreated cells or cells pre-treated with the unlabeled 6E competitor peptide. Membrane proteins often appear as indistinct bands when resolved with SDS-PAGE, consistent with our data. It is possible that binding of our crosslinking peptide to a cell surface protein facilitates labeling of other membrane proteins in proximity, as implicated by activation of receptor signaling *in trans* by tethered GPCR ligands in a previous study^34^. To visualize labeling, fluorescence microscopy was used to show the localization of **FAM-1(NO_2_)** upon labeling A2AR(Nb_6E_)-HEK cells (Figure 3F). As predicted, signal from **FAM-1(NO_2_)** and Nb_Alfa_ labeled with tetramethylrhodamine (TMR) co-localized at the cell periphery. Pretreatment with an unlabeled competitor peptide (6E) eliminated FAM but not TMR signal, as expected since **FAM-1(NO_2_)** and Nb_Alfa_ have different sites of binding on A2AR(Nb_6E_). Collectively these data demonstrate specific and rapid covalent labeling of living cells.

We then applied our PIR methodology to convert a non-covalent small molecule agonist of A2AR (CGS21680)^35^ into a covalent ligand through linkage with a crosslinking peptide. CGS21680 is a potent agonist of A2AR^36^. This compound contains a carboxylate group that projects from the extracellular vestibule of A2AR in a crystal structure^37^ and offers a convenient point for linkage to other moieties without substantial losses in biological activity^38^. To allow for site-specific linkage of the crosslinking peptide to CGS21680 we appended azide and alkyne moieties, respectively (Figure 4A). Using copper-catalyzed click chemistry, we prepared a **CGS-1(NO_2_)** conjugate as well as a non-reactive analog (**CGS-6E-A5**) that should have similar affinity for Nb_6E_ yet would not crosslink to the receptor (Figure 4B, Supporting Figure 11). We first evaluated these conjugates for the activation of signaling through A2AR(Nb_6E_) in HEK cells in conventional dose-response assays to monitor the production of the common GPCR second messenger molecule cyclic adenosine monophosphate (cAMP). A well-established cAMP-responsive luciferase variant^39^ was stably expressed in cells to quantitate ligand-induced receptor activation^40^. CGS21680 was slightly more potent than **CGS-6E-A5** and **CGS-1(NO_2_)** (Figure 4C, Supporting Figure 12) although all compounds were similarly efficacious at high doses. We also evaluated the kinetics of signal cessation following ligand removal (washout assay, Figure 4D, Supporting Figure 12). In this assay, **CGS-1(NO_2_)** cAMP responses persisted following ligand washout, whereas **CGS-6E-A5** and CGS21680 responses returned to baseline levels over the course of 30 minutes. It is likely that irreversible association of **CGS-1(NO_2_)** with the receptor via covalent binding explains these results. This finding leads us to hypothesize that ligand dissociation plays a larger role than cell intrinsic mechanisms of receptor signaling desensitization (β-arrestin recruitment and receptor internalization) in the context of this assay system and the ligands tested. This observation is in line with a previous study that applied an A2AR agonist that directly crosslinked to the receptor^41^. The ability of A2AR to induce protracted signaling may relate to the unusual propensity of this receptor to resist internalization upon activation^42^

**Figure 4.**
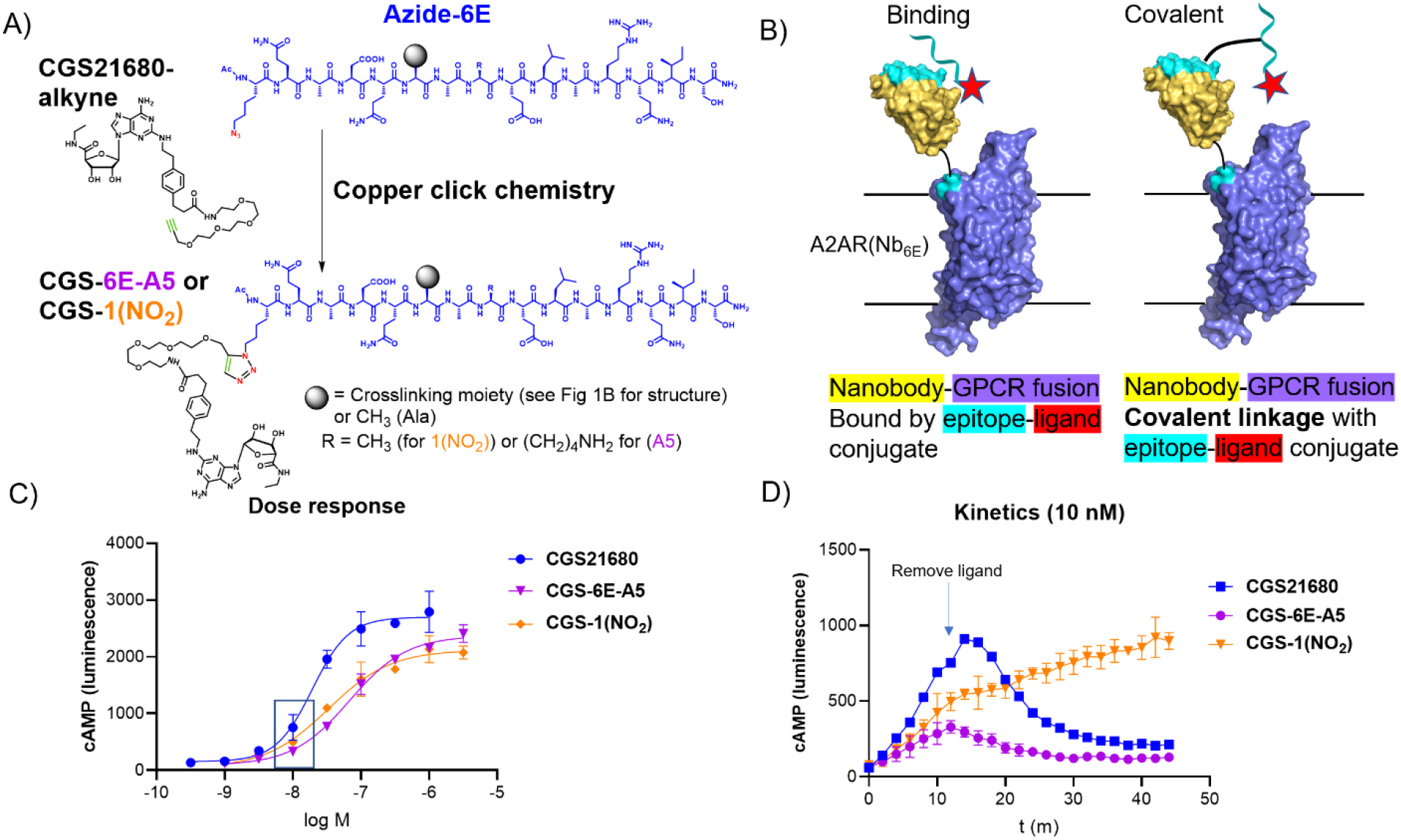
Assessment of biological consequences of epitope tag-mediated crosslinking on signaling properties. A) Synthetic scheme for synthesis of conjugates consisting of a 6E peptide analogue with or without crosslinking capacity and an A2AR ligand (CGS21680) using copper catalyzed azide alkyne cycloaddition chemistry. B) Schematic comparing association between a crosslinking peptide-ligand conjugate and a non-crosslinking peptide-ligand conjugate. C) Dose-response assays for the induction of cAMP responses in cells expressing A2AR(Nb_6E_). Curves are generated from a sigmoidal dose-response model with variable slope. Replicate data and tabulation of results are shown in Supporting Figures 12-13. D) Analysis of kinetics of ligand-derived signaling following the removal of free ligand (at 12 m) in a washout assay. For panels C-D, the data points show mean ± SD from a single experiment.

## Discussion

In this manuscript, we have described the first example of a nanobody-epitope tag pair that uses PIR to enable rapid and specific covalent bond formation. Other examples of antibodies and nanobodies that engage in PIR are challenging to apply in systems of choice because the epitopes are ill-defined or exist as part of a folded (i.e. non-linear) epitope^13–15,19,43^. In contrast, the system described here can be applied to different targets because the crosslinking epitope recognized is small, linear, readily amenable to synthesis, and can be linked to other synthetic compounds using click chemistry. Another beneficial feature of our platform is that the recognition element (Nb_6E_) can be linked to biological targets of choice through genetic fusion. The use of Nbs in place of conventional single-chain fragment variable (ScFv)-fusions can mitigate common issues such as construct aggregation and heavy chain-light chain domain mispairing^44^.

We also demonstrated that the fusion of Nb_6E_ with either of two GPCRs derived from different classes (A2AR class A and PTHR1 class B, Figure 3 and Supporting Figure 8), provides correctly folded receptors that can be covalently labeled with a crosslinking peptide. Further, both A2AR- and PTHR1-Nb_6E_ fusions effectively respond to conventional agonists (Figure 4, Supporting Figure 9). These successes suggest that this approach may be widely applicable, both to the large and biomedically important GPCR superfamily and potentially to other classes of membrane proteins. With expanding interest in identifying molecules that covalently target proteins of interest, a generalizable system for interrogating the impact of covalent ligation will be useful. This approach might also be useful for new types of bitopic ligands^45^.

Another valuable property of our platform is the rapid rate of labeling achieved through PIR. We have shown that covalent labeling of receptors on the surface of live cells occurs on the order of a few minutes, which is faster than other antibody/nanobody-PIR systems^13–15,19,43^ and is comparable to the rates of other enzymatic self-labeling domain tags (Halo- or SNAP-tags) and the fastest biorthogonal labeling chemistries^29–31^. Rapid labeling is essential for dissecting temporal features of proximal signaling responses of cell surface proteins^46^, which often peak a few minutes after stimulation and dissipate within an hour.

Our system differs from other self-labeling domain tags (SNAP, Halo) in that the recognition element (Nb_6E_) is adapted from a protein (alpaca heavy chain only antibody) naturally found at the cell surface in mammalian cells^24^. When prokaryotic or intracellular protein domains, such as SNAP and Halo, are applied as cell surface labels in cell engineering projects, there is a substantial possibility that immunogenic responses will be induced^47^. Nanobodies are generally weakly immunogenic, they have been applied as recognition elements in chimeric antigen receptor (CAR) T cells^48^, and one has been approved for therapeutic application in humans^24^. The data shown here suggest the covalent nanobody-epitope platform could be useful in the selective labeling of cells engineered to express designer receptors, such as CAR-T cells. Applications that require rapid labeling, such as those using short lived positron emitting (PET) isotopes, might also benefit from this platform.

## Acknowledgments

We acknowledge Daniel Appella, Carole Bewley, Samuel Gellman, and Kenneth Jacobson for helpful discussions. This work was supported by the NIH Intramural Research (NIDDK).

## Methods

### Synthesis

Peptides were synthesized using solid-phase peptide synthesis with Fmoc protection of backbone amines. Crosslinking groups were attached using cysteine-maleimide chemistry. All compounds were purified with reverse-phase HPLC, and compound identity was confirmed by mass spectrometry. Synthetic details are described in the supporting methods section. Mass spectrometry characterization is summarized in Supporting Table 1.

### Cell culture and cell-based assays

Cell-based experiments were run with clonal cell lines derived from HEK293 (ATCC CRL-1573) stably transfected with a biosensor for cAMP^39^ and receptors of interest^40^. Receptor sequences are found in Supporting Methods. Biological response assays were run in white walled 96-well plates and luminescence responses were recorded on a plate reader, as described previously^40^ and in Supporting Methods. Cell-based binding assays were performed with freshly trypsinized cells and analysis was performed using flow cytometry as described in Supporting Methods.

### Protein expression and labeling

Nanobodies were expressed in the *E. coli* periplasm and purified as previously described^26^ and detailed in Supporting Methods. The sequence of Nb_6E_ has been published previously^25,26^. Purified nanobodies were site specifically labeled using sortagging as previously described^49^ and detailed in the Supporting Methods.

### Analysis of peptide crosslinking

Peptide crosslinking was analyzed by several complementary methods including mass spectrometry, flow cytometry, SDS-PAGE, and Western blotting. LC/MS quantitation of nanobody-peptide crosslinking was performed on a Waters Xevo qTof LC/MS as described in Supporting Methods. SDS-PAGE was run using 15% acrylamide gels prepared in house and gel-based fluorescence measurements were made on a gel imager as described in Supporting Methods. Western blotting analysis of cell surface protein labeling was performed using a fluorescein-specific monoclonal antibody conjugated to horse radish peroxidase (Supporting Methods).

### Data calculations and display

Data were analyzed and prepared for display using GraphPad Prism, FlowJo, and Fiji (ImageJ). Flow cytometry data were quantified by measuring median fluorescence of intensity (MFI) measurements. Kinetic rate constants were calculated as described previously^30^ and in Supporting Methods.

### Fluorescence Microscopy

HEK293 cells were stained, fixed, and analyzed as described in Supporting Methods.

### Assessment of Nb-peptide crosslinking site using MS/proteomics

Crosslinked Ac-1-Nb_6E_ was digested with proteases and analyzed using mass spectrometry to identify nanobody residues involved in crosslinking. We searched for fragments resulting from crosslinking at Lys, His, Ser, and Thr residues, based on literature precedent^50^. Methodology is described in Supporting Methods.

## Notes

### Competing Interest Statement

The authors have declared no competing interest.

