## Supporting Information for "Rapid covalent labeling of a GPCR on living cells using a nanobody-epitope tag pair to interrogate receptor pharmacology"

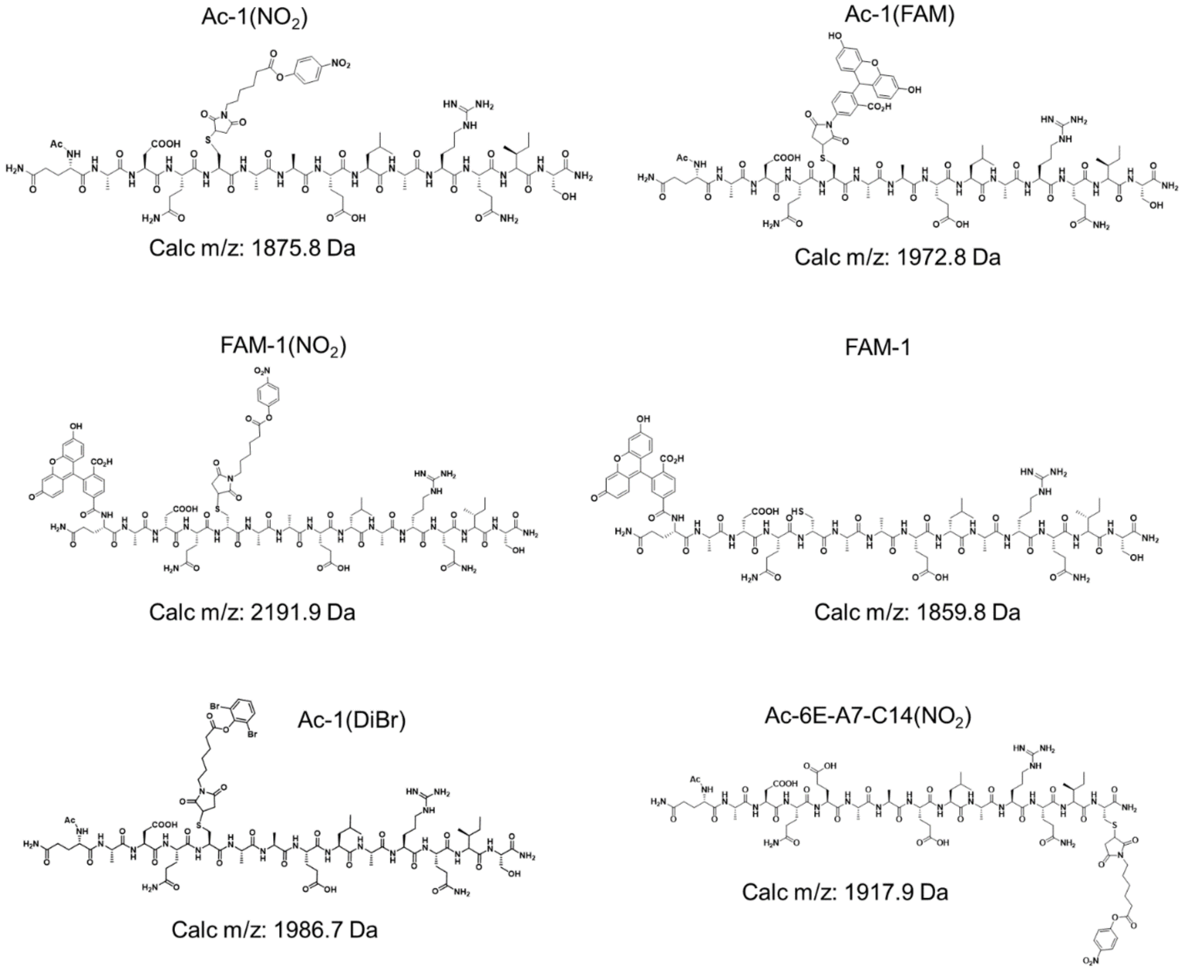


**Supporting Figure 1.** Structures and calculated m/z of various peptides used in this study. See Supporting Table 1 for a summary of experimental mass spectrometry data.

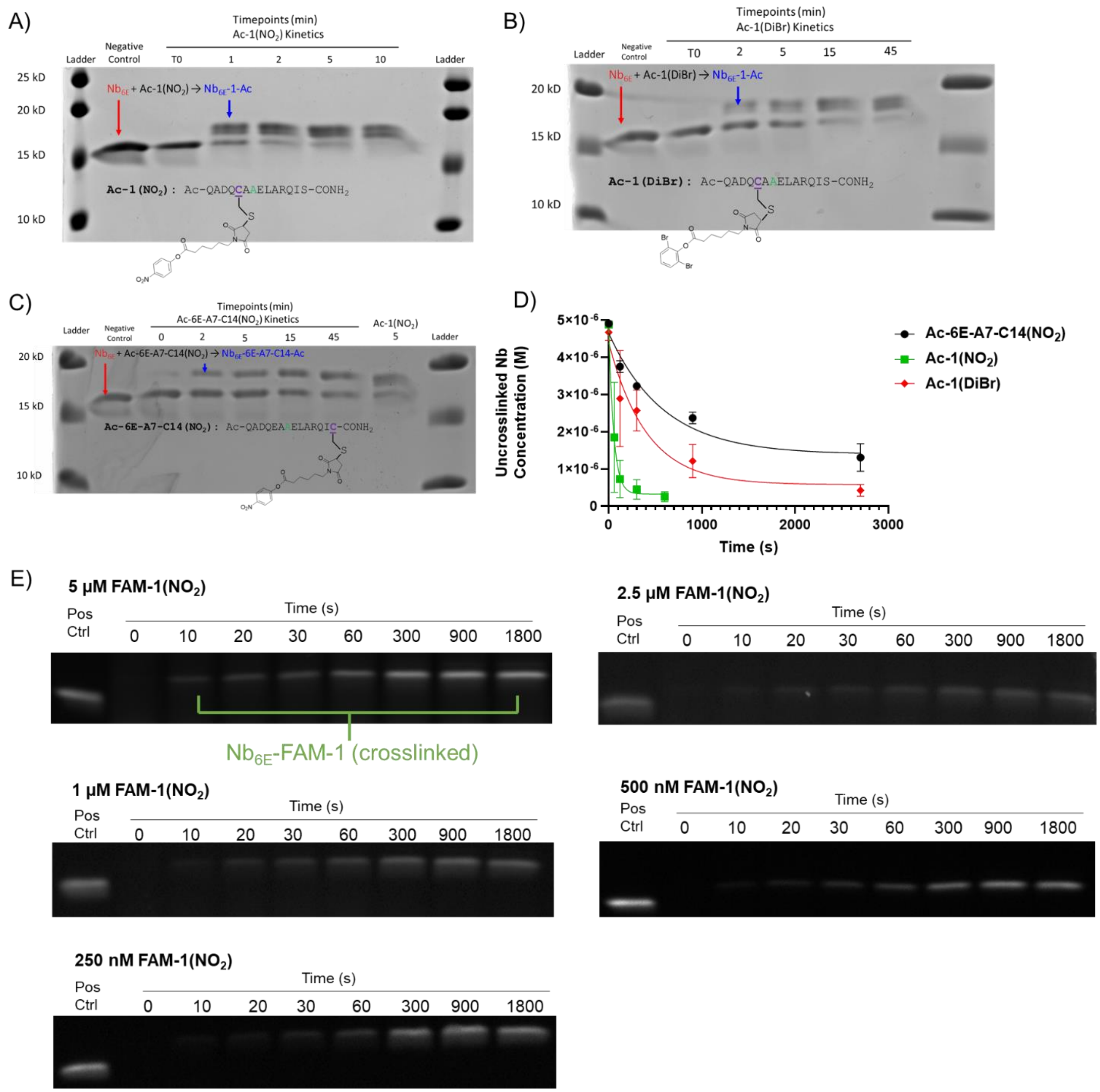


**Supporting Figure 2.** Assessment of crosslinking kinetics. SDS-PAGE showing time-course crosslinking between $Nb_{6E}$ and 6E variants A) Ac-1($NO_2$), B) Ac-1(DiBr), and C) Ac-6E-A7-C14($NO_2$). Crosslinked product appears as a higher molecular weight band than starting material. D) Assessment of crosslinking kinetics using LC/MS. The abundance of uncrosslinked nanobody was quantified as described in Methods. Data points and error bars correspond to mean ± SD from three independent experiments. Pseudo first order rate constants are reported in the main text. E) Time course in-gel fluorescence of $Nb_{6E}$ (100 nM) crosslinked with indicated concentrations of Ac-1($NO_2$). Samples were resolved by SDS-PAGE and analyzed as described in Methods. "Pos Ctrl" refers to a nanobody labeled at a single site with fluorescein prepared by sortagging. Data from these experiments were used to calculate the $K_D$ and second order rate constant reported in the main text.

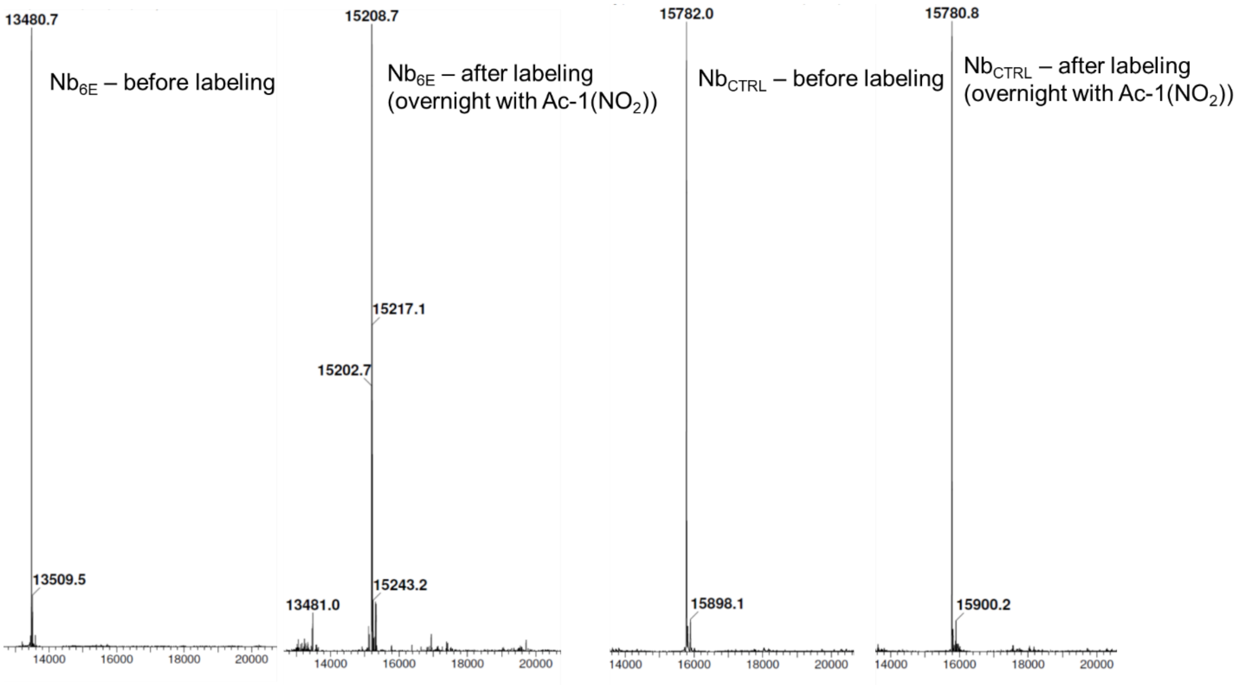


**Supporting Figure 3:** Crosslinking of Ac-1($NO_2$) is specific for $Nb_{6E}$. Indicated nanobodies (5 μM) were incubated with Ac-1($NO_2$) (20 μM) in PBS overnight at 4°C. Samples were analyzed by LC/MS as described in methods. $Nb_{CTRL}$ is a nanobody that binds to an irrelevant cell surface marker (CD38).

| Crosslinking Position | Residue identity | Score | PEP | Intensity |
|---|---|---|---|---|
| 7 | S | 236 | 2.41E-256 | 2.0E+08 |
| 17 | S | 355 | 0 | 2.6E+08 |
| 43 | K | 122 | 3.92E-40 | 1.3E+09 |
| 61 | S | 56 | 0.00014909 | 9.0E+07 |
| 63 | K | 134 | 1.14E-36 | 4.9E+09 |
| 69 | S | 155 | 2.07E-74 | 2.0E+07 |
| 74 | K | 114 | 3.51E-30 | 2.2E+06 |
| 85 | K | 112 | 1.61E-15 | 1.3E+07 |

**Supporting Figure 4.** Characterization of sites of peptide-nanobody crosslinking using mass spectrometry-proteomics. The crosslinking position is relative to the sequence with the pelB leader sequence removed (begins QVQ). “Score” corresponds to the Andromeda search engine within MaxQuant assigning confidence to the identity fragment assigned. “PEP” corresponds to the Bayesian posterior error probability from the search. “Intensity” corresponds to the intensity of the observed ions.

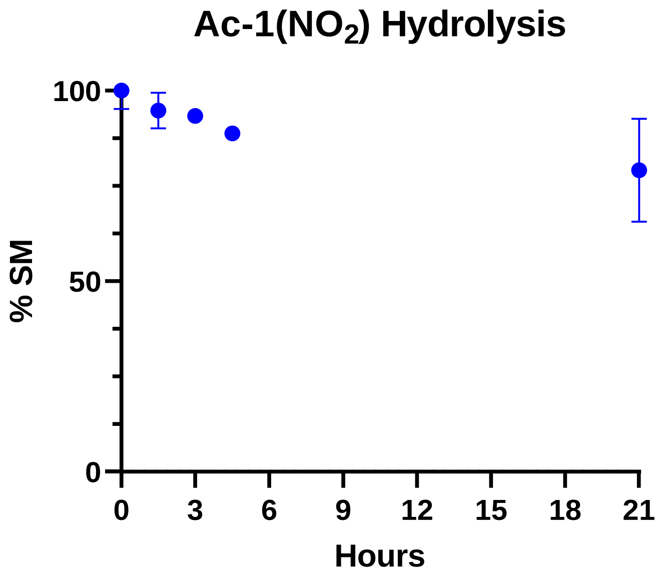


**Supporting Figure 5.** Time course analysis of hydrolysis of Ac-1($NO_2$) in PBS. % Starting material (SM) was monitored by analytical reverse-phase HPLC. Points represent duplicate runs normalized to the average of areas under the peak observed in HPLC traces.

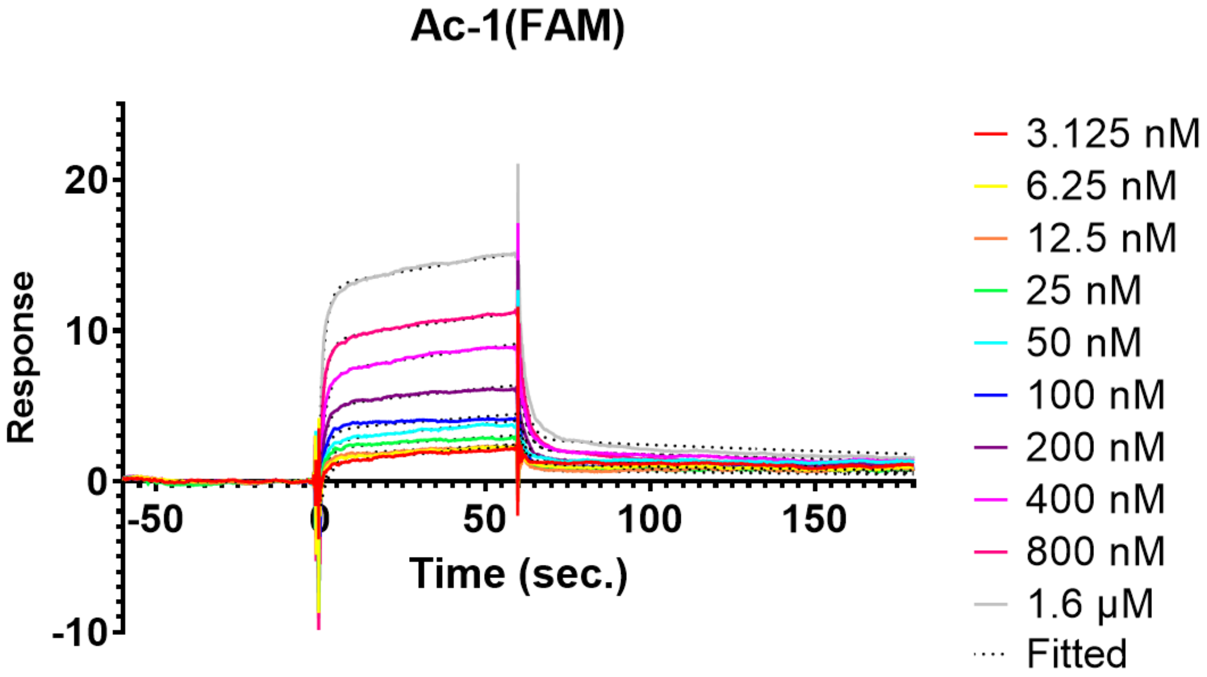


| Peptide Analyte | Mean $K_D$ (nM) | SD (nM) | n |
|---|---|---|---|
| Ac-1(FAM) | 1160 | 1000 | 4 |

**Supporting Figure 6**. (Top) Representative surface plasmon resonance sensorgram of Ac-1(FAM) binding to immobilized $Nb_{6E}$-biotin. Colored lines represent data points and dotted lines represent application of the binding model. See Supporting Figure 1 for structure. The experiment was performed as described in methods. (Bottom) Tabulation of sensorgram results from four independent experiments.

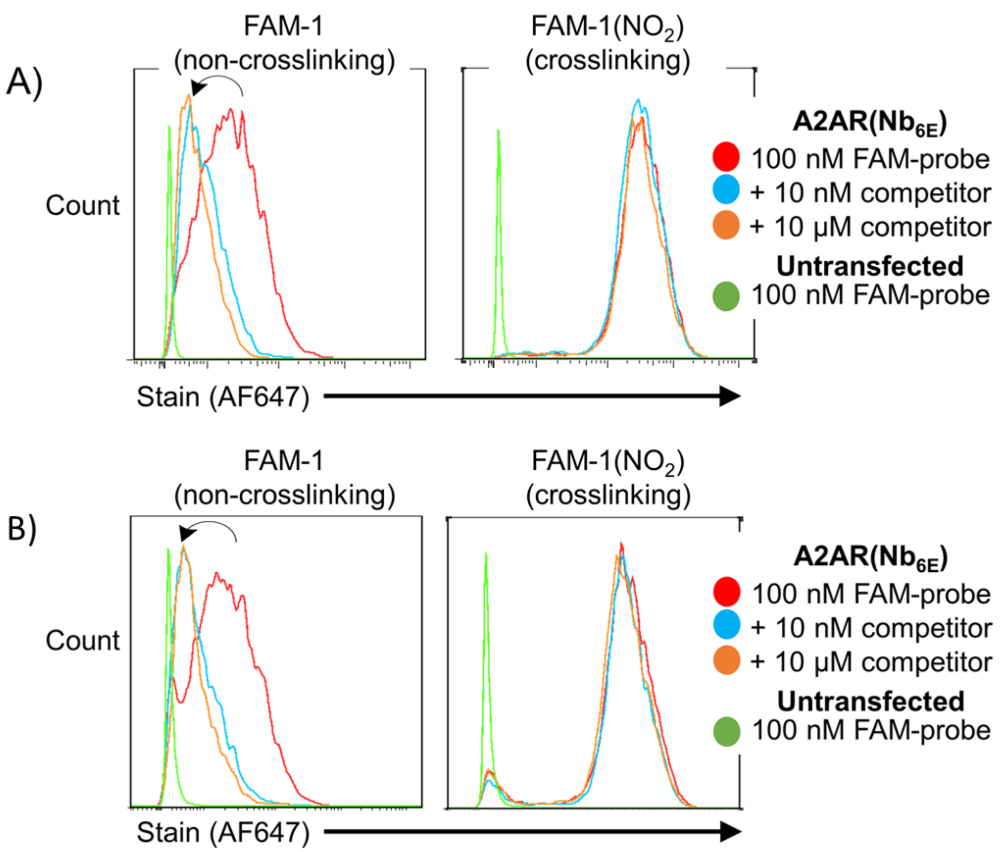


**Supporting Figure 7.** Independent replicates for evaluation of FAM-1 and FAM-1($NO_2$) labeling HEK293 cells expressing A2AR($Nb_{6E}$). Panels A and B correspond to independent replicate experiments. Cells were prepared and stained for analysis as described in Methods. Data are presented as histograms with staining in the Alexafluor647 channel on the X-axis. Green traces depict the same labeling experiment run cells untransfected cells that do not express the engineered receptor construct.

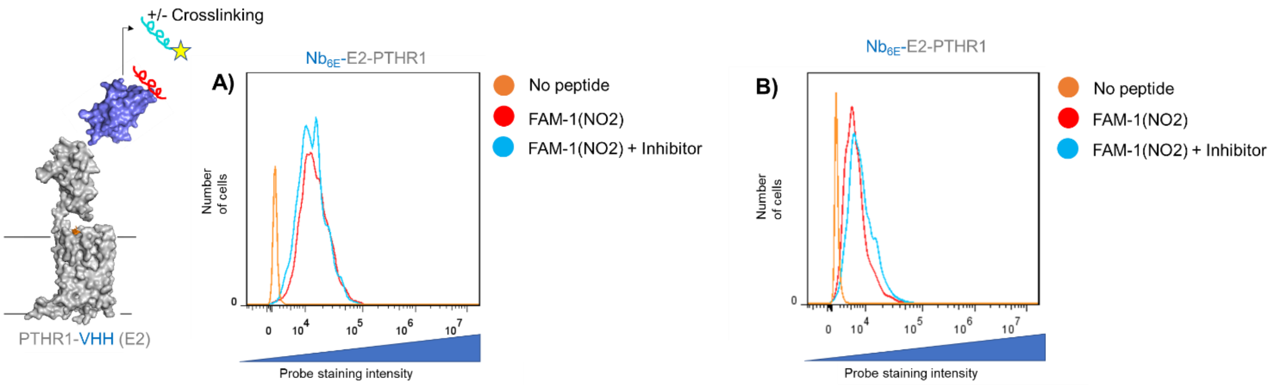


**Supporting Figure 8.** Analysis of HEK293 cells expressing $Nb_{6E}$-E2-PTHR1. Note that this is a different receptor than that shown in Figure 3. Cells were incubated with FAM-1($NO_2$) (100 nM) with or without the subsequent addition of a high concentration of unlabeled competitor peptide (“inhibitor” refers to 6E peptide, 10 μM). No peptide refers to cells not exposed to FAM-labeled peptide. Sequence information for this receptor is presented in Supporting Methods. Data are presented as histograms with staining in the Alexafluor647 channel on the X-axis. Each histogram corresponds to an independent experiment.

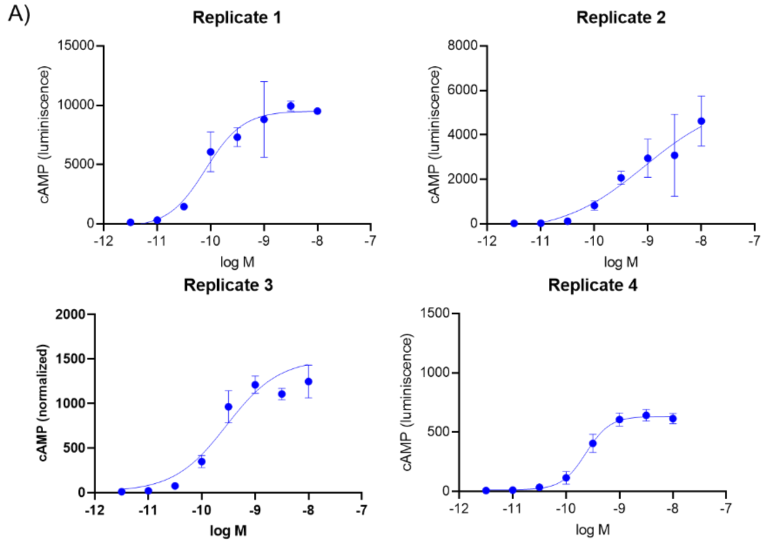


B)

| | $EC_{50}$ (nM ± SD) | | |
|---|---|---|---|
| PTH1-34 | 0.37 | ± | 0.35 |

Vs 0.51 nM for WT hPTHR1 in Cheloha Nature Comm 2020

**Supporting Figure 9**. Assessment of activation of $Nb_{6E}$-E2-PTHR1 with PTH(1-34). A) Data points represent mean ± SD from technical replicates in a single experiment. Curves correspond to fitting of a 4-parameter sigmoidal dose-response model. B) Tabulation of receptor activation parameters from four independent experiments. Cells were stably transfected with receptor and cAMP biosensor and were treated with PTH1-34 at indicated concentrations. Receptor activation was measured as described in Methods.

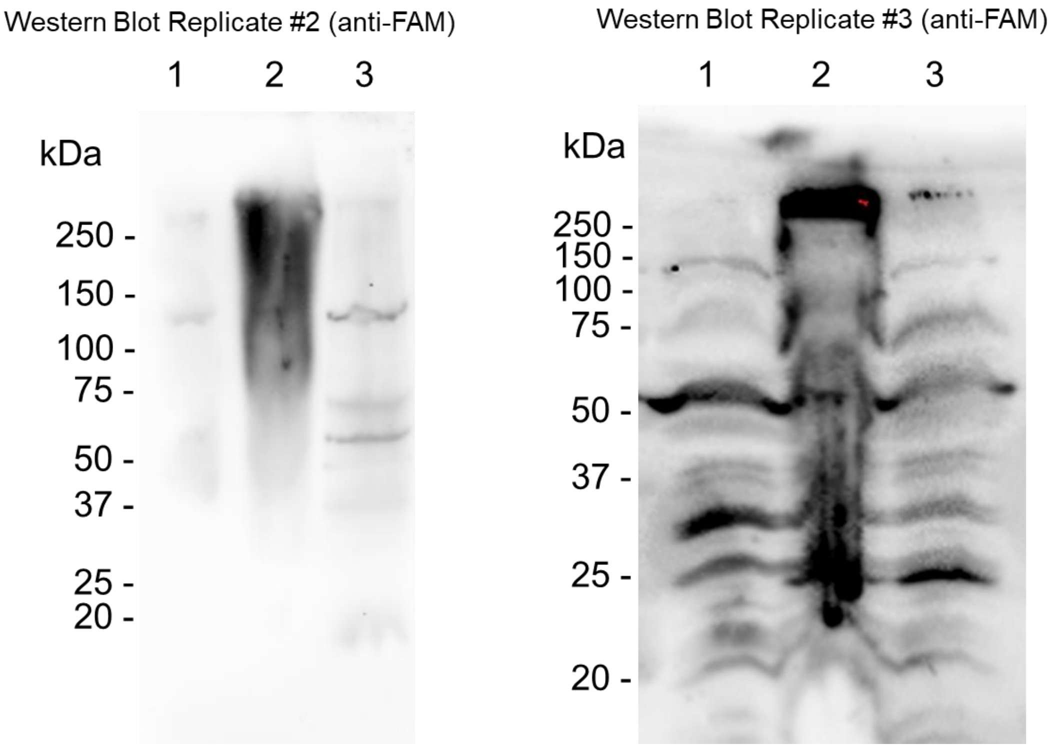


**Supporting Figure 10.** Replicate Western Blots of crosslinking with Fam-1($NO_2$). Lane 1: No probe added; Lane 2: 100 nM FAM-1($NO_2$); Lane 3: Pretreatment with 10 μM unlabeled competitor peptide (6E) followed by treatment with 100 nM FAM-1($NO_2$). Western Blotting was performed as described in Supporting Methods.

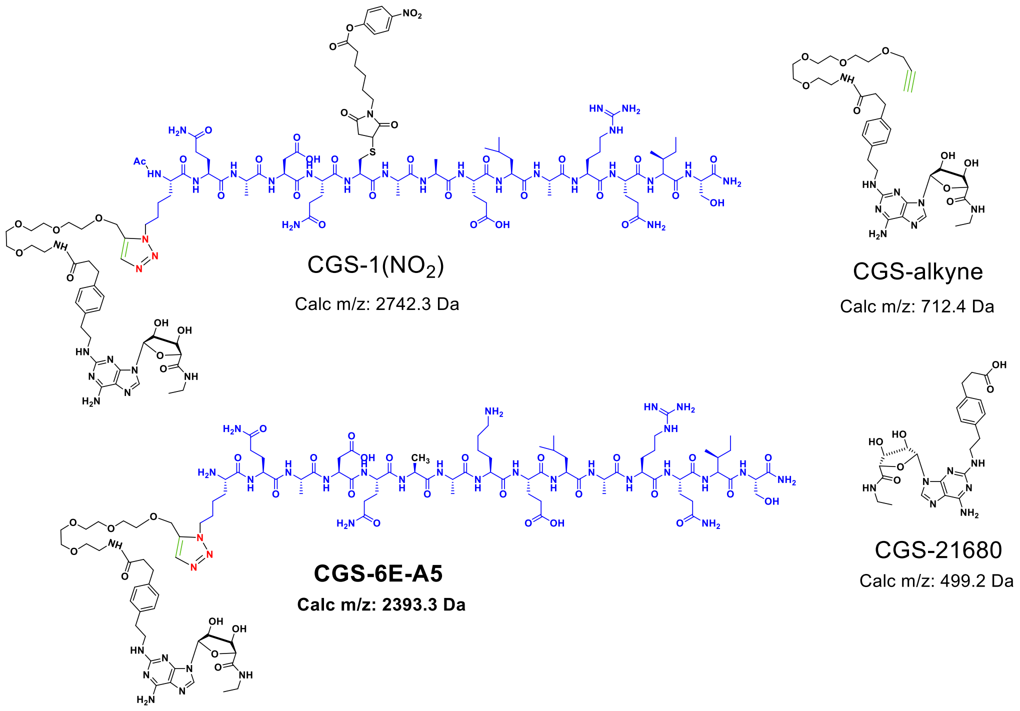


**Supporting Figure 11.** Structures and calculated m/z of CGS-1($NO_2$) and its noncrosslinking analog CGS-6E-A5. Compounds were synthesized as described in the methods section. These compounds were characterized by mass spectrometry (See Supporting Table 1)

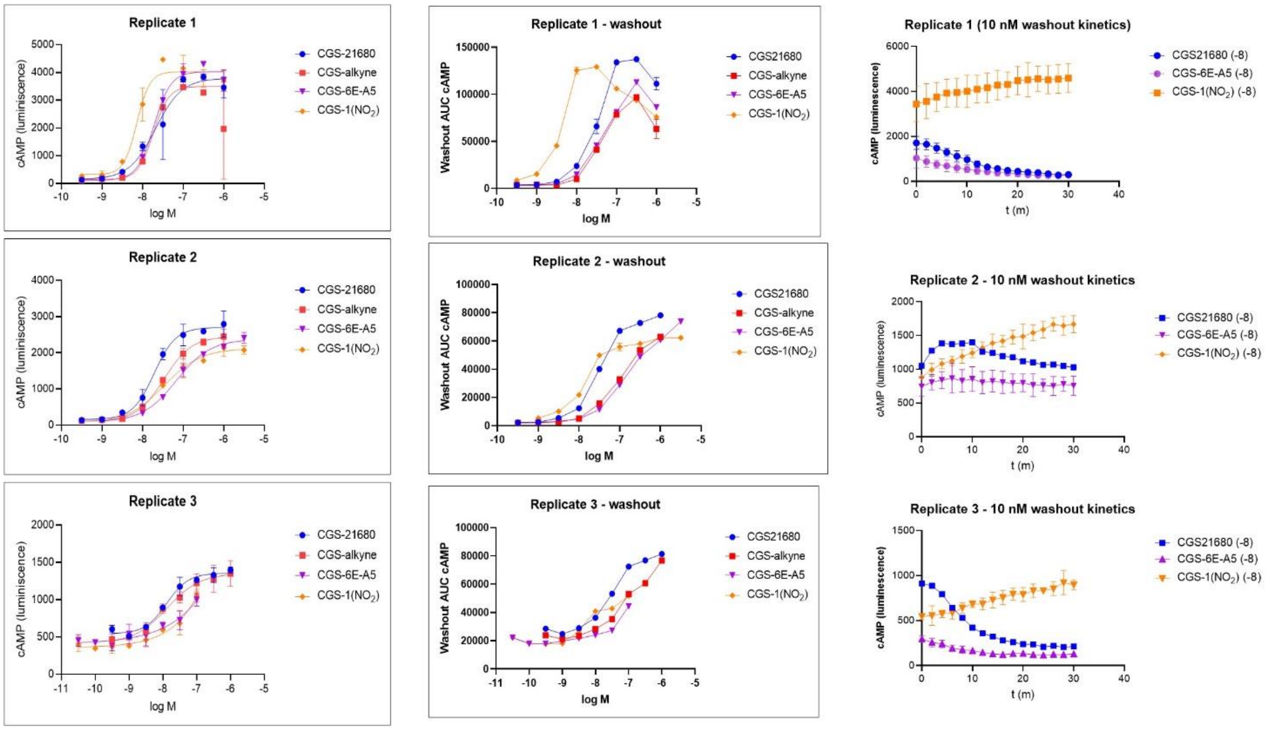


**Supporting Figure 12.** Replicate data for activation of A2AR(Nb$_{6E}$) signaling in HEK293 cells by CGS-6E variant conjugates. Data sets on the left correspond to dose-response curves for maximal cAMP responses generated upon addition of ligands to cells for 12 minutes. Curves were generated by fitting data to a 4-parameter sigmoidal dose-response model. Data sets in the middle column correspond to values generated from quantifying the area under the curve for kinetic washout responses upon removing ligand and measuring cAMP signaling ("washout") for the next 30 minutes. Lines connecting data points are shown to guide the eye. Data sets on the right correspond to kinetic plots for cAMP responses following washout for 10 nM doses of indicated peptides. Replicate experiment number 3 in this figure is the same as main Figure 4C-D.

| | Ligand on | | Washout | | Ratio | |
|---|---|---|---|---|---|---|
| | $EC_{50}$, nM | SD | AUC $EC_{50}$, nM | SD | (Ligand on/washout $EC_{50}$) | n |
| CGS-21680 | 15 | 5 | 27 | 2 | 1.8 | 3 |
| CGS-alkyne | 23 | 12 | 118 | 21 | 5.1 | 3 |
| CGS-6E-A5 | 57 | 34 | 125 | 64 | 2.2 | 3 |
| CGS-6E-C5(NO2) | 47 | 36 | 35 | 35 | 0.8 | 3 |

**Supporting Figure 13**. Tabulation of biological activity of A2AR ligand conjugates. $EC_{50}$ values correspond to mean measurements from three independent experiments. "Ligand on" data sets are prepared from the maximal signal recorded after ligand addition. "Washout" data are derived from integrated area under the curve values from kinetic measurements of cAMP/luminescence following ligand removal. See Figure 12 for graphical depictions of data. Compound structures are shown in Supporting Figure 1.

**Supporting Table 1**

| **Compound** | **Calc m/z (Da)** | **Observed m/z (Da)** |
|---|---|---|
| Ac-1($NO_2$) | 1875.8 | 1875.8 |
| Ac-1(FAM) | 1972.8 | 1971.8 |
| FAM-1 | 1859.8 | 1859.8 |
| FAM-1($NO_2$) | 2191.9 | 2193 |
| Ac-1(DiBr) | 1986.7 | 1988.6 |
| Ac-6E-A7-C14($NO_2$) | 1917.9 | 1917.8 |
| CGS-Alkyne | 712.4 | 712.3 |
| Azide-6E-A5 | 1680.9 | 1681.6 |
| CGS-6E-A5 | 2393.3 | 2392.2 |
| Azide-1 | 1697.8 | 1697.6 |
| Azide-1($NO_2$) | 2029.9 | 2030.6 |
| CGS-1($NO_2$) | 2742.3 | 2741.8 |

**Supporting Methods.**

**Data analysis.** Average values and uncertainties correspond to mean ± standard deviation. Dose-response data were fit to 4-parameter sigmoidal dose-response model. For data sets where the top plateau was not reached, models were constrained to match the top plateau value reached by full agonists.

**Cell culture and cell lines.** HEK293 cells (ATCC CRL-1573) were stably transfected with a cAMP responsive luciferase variant[1] and the receptor of interest to provide clonal cell lines that could be grown without selection antibiotic. All cell lines were cultured in DMEM supplemented with 10% fetal bovine serum and penicillin/streptomycin. Cells were checked for mycoplasma contamination.

**Measurement of receptor activation using cAMP-responsive luciferase expression.** Assays were run according to previously described protocols[2]. Briefly, cells were plated in a white walled, clear bottom 96 well plate (Corning #3610) and grown to confluency. Growth medium was removed and $CO_2$ independent medium containing luciferin (0.5 mM) was added until a stable background reading was obtained (~10 minutes). 10x stocks of ligands were then added to wells and cAMP (luminescence) responses were measured for every 2 minutes for 12 minutes after ligand addition (Biotek Neo2 plate reader). The response recorded 12 minutes after ligand addition was used for construction of dose-response curves.

**Receptor signaling (cAMP) washout assay.** cAMP responses induced by ligand addition were measured as above. After 12 minutes the medium containing ligands was discarded. New $CO_2$ independent medium containing fresh luciferin was added to all wells and luminescence responses were measured every 2 minutes for an additional 30 minutes as described above. Area under the curve (AUC) values were used to construct dose-response curves for washout assays.

**Protein expression plasmids.** Nanobodies were expressed in *E. coli* from pET26b(+) plasmids encoding a sequence corresponding to pelB leader-nanobody-LPETGG-His6 under the control of Lac repressor (custom cloning from GenScript). The sequence of VHH05/VHH-6E/$Nb_{6E}$ has been published previously[2,3].

**Receptor Plasmids.** Custom plasmids were ordered from Vectorbuilder. Plasmids included elements encoding G418 resistance and a CAG promoter for the protein of interest. Insert sequences are listed below here.

HGH secretion tag-alfa tag-$Nb_{6E}$-human A2AR

MATGSRTSLLLAFGLLCLPWLQEGSAFPTIPLSGSEPSRLEEELRRRLTEPSGSMAQVQLQES
GGGLVQPGGSLRLSCAASGFVFENSAMAWYRQAPGKERELIAVIGTTFIKLAESVKGRFTISRD
NAKSTVYLQMNNLKPEDTAVYYCSKSGAYWGQGTQVTVSSGGLPETGGSGGMPIMGSSVYIT
VELAIAVLAILGNVLVCWAVWLNSNLQNVTNYFVVSLAAADIAVGVLAIPFAITISTGFCAACHGC
LFIACFVLVLTQSSIFSLLAIAIDRYIAIRIPLRYNGLVTGTRAKGIIAICWVLSFAIGLTPMLGWNNC
GQPKEGKNHSQGCGEGQVACLFEDVVPMNYMVYFNFFACVLVPLLLMLGVYLRIFLAARRQL
KQMESQPLPGERARSTLQKEVHAAKSLAIIVGLFALCWLPLHIINCFTFFCPDCSHAPLWLMYLA
IVLSHTNSVVNPFIYAYRIREFRQTFRKIIRSHVLRQQEPFKAAGTSARVLAAHGSDGEQVSLRL
NGHPPGVWANGSAPHPERRPNGYALGLVSGGSAQESQGNTGLPDVELLSHELKGVCPEPPG
LDDPLAQDGAGVS*

atggcaacaggatcaaggacatccttgcttctcgcattcggccttctctgcctgccttggctgcaagagggtagcgcatttcctaccatac
ccttgtccggaagtgaaccctctaggctggaggaggaattgagacgccggttgacagagccctccggatccatggcacaggtccag
cttcaagagtccggtggtggtctggtgcagcccggcggttcactgcgccttagctgtgcggcaagcggctttgtttttgaaaacagcgca
atggcgtggtaccggcaggccccggggaaagagcgggagctcattgctgtgattggcactacttttattaaacttgcagaaagcgtta

agggccggtttacaatcagcagggataacgcgaagagcacagtatatctgcaaatgaacaacttgaagcctgaggacactgcagt
ctactattgtagtaagtctggtgcgtattggggacagggtacacaggtgaccgtcagctctggtggtcttcccgaaaccgggggttccgg
gggaatgccaatcatggggagctctgtatatatcacggtcgaactcgccattgctgtactggctattttgggcaacgttttggtttgctggg
cggtatggcttaatagcaatcttcaaaatgtgactaattacttcgtggtctcccttgctgcagccgatatcgccgttggtgttttggcgatacc
attcgcgatcaccatctctaccggcttttgtgctgcctgccatggctgcttgttcattgcttgttttgttttggtgctcacgcaatccagtatcttca
gtctcctcgcaatcgctatagatagatatatagcgatacgaatcccattgaggtacaacggactcgtaacaggaacaagagcgaaa
ggcataatagcaatttgctgggtgctctccttcgctattggactgaccccaatgctgggatggaacaattgtgggcaaccaaaagagg
gaaaaaaccacagccagggttgcggagagggtcaagtcgcttgcctttttgaagacgttgtcccaatgaattacatggtatattttaatttt
tttgcgtgcgtattggtacctttgctgctcatgctcggagtctatcttagaatatttcttgcggctcgacgacaactgaagcaaatggagtca
cagcctttgcctggggagagggcaagatctacccttcagaaggaggtgcatgcagcaaaaagccttgccataatcgtaggcctgttc
gctctttgctggttgccacttcatatcatcaactgtttcacgttcttttgtccagattgcagtcatgcgccgttgtggttgatgtatctggcaatcg
tgctgtcacataccaattcagttgttaacccatttatctatgcctatcgcatccgggagtttcgccaaacttttaggaagataatcaggagtc
atgtcctgcggcagcaggaaccatttaaagcggcagggacatccgcacgcgttttggcggcgcatggatcagacggggagcaagt
atcattgcgcctcaacgggcacccccctggtgtttgggctaatggatcagccccgcacccagaacggcggcctaatggttacgccttg
ggcttggtgagcgggggctccgcccaggagtcacaggggaacactggccttcctgacgtggaactcttgagtcacgagctgaaggg
cgtttgcccagagcctcccggactcgatgatccactggctcaagatggggctggcgtgtcatga

Human PTHR1(N)-$Nb_{6E}$-Human PTHR1(C)

MGTARIAPGLALLLCCPVLSSAYALVDADDVMTKEEQIFLLHRAQAQCEKRLKEVLQRPASIME
SDKGWTSASTSGKPRKDKASGKLYPESEEDKGGGSGGGSMAQVQLQESGGGLVQPGGSLR
LSCAASGFVFENSAMAWYRQAPGKERELIAVIGTTFIKLAESVKGRFTISRDNAKSTVYLQMNN
LKPEDTAVYYCSKSGAYWGQGTQVTVSSGGLPETGGEAPTGSRYRGRPCLPEWDHILCWPL
GAPGEVVAVPCPDYIYDFNHKGHAYRRCDRNGSWELVPGHNRTWANYSECVKFLTNETRER
EVFDRLGMIYTVGYSVSLASLTVAVLILAYFRRLHCTRNYIHMHLFLSFMLRAVSIFVKDAVLYSG
ATLDEAERLTEEELRAIAQAPPPPATAAAGYAGCRVAVTFFLYFLATNYYWILVEGLYLHSLIFM
AFFSEKKYLWGFTVFGWGLPAVFVAVWVSVRATLANTGCWDLSSGNKKWIIQVPILASIVLNFIL
FINIVRVLATKLRETNAGRCDTRQQYRKLLKSTLVLMPLFGVHYIVFMATPYTEVSGTLWQVQM
HYEMLFNSFQGFFVAIIYCFCNGEVQAEIKKSWSRWTLALDFKRKARSGSSSYSYGPMVSHTS
VTNVGPRVGLGLPLSPRLLPTATTNGHPQLPGHAKPGTPALETLETTPPAMAAPKDDGFLNGS
CSGLDEEASGPERPPALLQEEWETVM*

atggcaacaggatcaaggacatccttgcttctcgcattcggccttctctgcctgccttggctgcaagagggtagcgcatttcctaccatac
ccttgtccggaagtgaaccctctaggctggaggaggaattgagacgccggttgacagagccctccggatccatggcacaggtccag
cttcaagagtccggtggtggtctggtgcagcccggcggttcactgcgccttagctgtgcggcaagcggctttgtttttgaaaacagcgca
atggcgtggtaccggcaggccccggggaaagagcgggagctcattgctgtgattggcactacttttattaaacttgcagaaagcgtta
agggccggtttacaatcagcagggataacgcgaagagcacagtatatctgcaaatgaacaacttgaagcctgaggacactgcagt
ctactattgtagtaagtctggtgcgtattggggacagggtacacaggtgaccgtcagctctggtggtcttcccgaaaccgggggttccgg
gggaatgccaatcatggggagctctgtatatatcacggtcgaactcgccattgctgtactggctattttgggcaacgttttggtttgctggg
cggtatggcttaatagcaatcttcaaaatgtgactaattacttcgtggtctcccttgctgcagccgatatcgccgttggtgttttggcgatacc
attcgcgatcaccatctctaccggcttttgtgctgcctgccatggctgcttgttcattgcttgttttgttttggtgctcacgcaatccagtatcttca
gtctcctcgcaatcgctatagatagatatatagcgatacgaatcccattgaggtacaacggactcgtaacaggaacaagagcgaaa
ggcataatagcaatttgctgggtgctctccttcgctattggactgaccccaatgctgggatggaacaattgtgggcaaccaaaagagg
gaaaaaaccacagccagggttgcggagagggtcaagtcgcttgcctttttgaagacgttgtcccaatgaattacatggtatattttaatttt
tttgcgtgcgtattggtacctttgctgctcatgctcggagtctatcttagaatatttcttgcggctcgacgacaactgaagcaaatggagtca
cagcctttgcctggggagagggcaagatctacccttcagaaggaggtgcatgcagcaaaaagccttgccataatcgtaggcctgttc
gctctttgctggttgccacttcatatcatcaactgtttcacgttcttttgtccagattgcagtcatgcgccgttgtggttgatgtatctggcaatcg
tgctgtcacataccaattcagttgttaacccatttatctatgcctatcgcatccgggagtttcgccaaacttttaggaagataatcaggagtc

atgtcctgcggcagcaggaaccatttaaagcggcagggacatccgcacgcgttttggcggcgcatggatcagacggggagcaagt
atcattgcgcctcaacgggcacccccctggtgtttgggctaatggatcagccccgcacccagaacggcggcctaatggttacgccttg
ggcttggtgagcgggggctccgcccaggagtcacaggggaacactggccttcctgacgtggaactcttgagtcacgagctgaaggg
cgtttgcccagagcctcccggactcgatgatccactggctcaagatggggctggcgtgtcatga

**Nanobody Purification from *E. coli.*** BL21(DE3) *E. coli* were transfected via heat shock with pET26b(+) plasmids encoding nanobodies of interest and grown in medium (Terrific Broth) containing kanamycin (50 µg/mL). Transformed bacteria were used generate a starter culture, which was used to inoculate full-size cultures (1-4 L) containing kanamycin. This culture was grown at 37°C with growth monitored through measurement of the optical density at 600 nm ($OD_{600}$). When $OD_{600}$ values between 0.3 and 0.8 were observed, protein expression was induced by addition of Isopropyl β-d-1-thiogalactopyranoside (IPTG, 1 mM). The induced culture was then shaken 30°C overnight.

Bacteria were harvested via centrifugation for 30 min at 6,000 RPM (Avanti J Series centrifuge). Cells were resuspended in 30 mL of NTA wash buffer (tris buffered saline + 10 mM imidazole, pH 7.5) containing protease inhibitor (Pierce Protease Inhibitor Tablets, ThermoFisher A32953). Cells were then lysed via sonication and the lysate was centrifuged at 15,000 RPM for 45 min. The supernatant was then passed through a fritted column containing nickel NTA beads (His Pur™ Ni-NTA Resin) equilibrated with Nickel NTA wash buffer. After initial flowthrough, beads were washed 3x with Nickel NTA wash buffer. Subsequently, bound protein of interest was eluted using 10 mL of Nickel NTA elution buffer (TBS + 150 mM imidazole, pH 7.5). Sample was subjected to size exclusion chromatography (Cytiva Akta™ / Pure) using a HiLoad ™ 16/600 Superdex 200 pg column with an isocratic gradient of TBS (Flow rate 1 mL/min). Fractions of interest were collected and concentrated via centrifugation using spin filtration columns (Amicon Ultra-15, regenerated cellulose, 10,000 nominal molecular weight limit). Protein concentrations were determined absorption at 280 nm.

**Peptide Synthesis, Cleavage, and Purification.** All peptides were synthesized via Fmoc solid phase peptide synthesis on a Gyros PurePep Chorus Automated Peptide Synthesizer. Peptide assembly was performed on Rink Amide resin (0.05 mmol scale) to afford a C-terminal carboxamide. Fmoc-amino acids were dissolved in dimethylformamide (DMF) and added to resin (8 equivalents) with HATU ((1-[Bis(dimethylamino)methylene]-1H-1,2,3-triazolo[4,5-b]pyridinium 3-oxid hexafluorophosphate, 8 equivalents) and N,N-diisopropylethylamine (DIPEA, 16 equivalents). Fmoc groups were deprotected using 20% piperidine in DMF. For peptides bearing a N-terminal fluorescein, the fluorophore was manually coupled after automated linear synthesis using 10 equivalents of 5(6)-Carboxyfluorescein (Acros Organics), HATU (10 eq), and DIPEA (20 eq) and overnight incubation at room temperature.

Cleavage of peptides was performed using a cleavage cocktail comprised of trifluoroacetic acid(TFA)/$H_2O$/triisopropylsilane(TIS) (92.5:5:2.5% by volume). Variants containing cysteine residues were cleaved using the following cleavage cocktail TFA/$H_2O$/TIS/ethanedithiol (EDT) (90:5:2.5:2.5 by volume) and rocked at room temperature for 3 hours prior to filtration. Product was precipitated using chilled diethyl ether and pelleted by centrifugation (3,000 RPM for 2 minutes). Diethyl ether was decanted, and the pellet was dried under $N_2$ prior to being dissolved in DMSO and purified by HPLC. Peptides were purified via preparative-scale HPLC using a Phenomenex Aeris Peptide XB-C18 Prep column (particle size 5 µM, 100 Å pore size) with a linear gradient of solvent A (0.1% TFA in $H_2O$) and solvent B (0.1% TFA in acetonitrile). Fractions of interest were combined and lyophilized. Lyophilized peptides are then dissolved in DMSO at desired concentrations and frozen.

**Peptide Crosslinking assessment by SDS-PAGE.** Crosslinking reactions were performed at room temperature with a final concentration of 20 µM of crosslinking 6E peptide and 5 µM $Nb_{6E}$ in PBS. At predetermined time points, 20 µL of crosslinking reaction was quenched by addition of 5 µL ethanolamine (10 M). Prequenched samples ($t_0$) were generated by mixing $Nb_{6E}$ and ethanolamine prior to addition of crosslinking peptide. All samples were neutralized with $H_2SO_4$ (6M) prior to analysis. A portion of each quenched sample was reserved for mass spectrometry (see below), and the rest used for analysis by SDS-PAGE. Briefly, each sample was denatured with SDS-PAGE loading buffer containing 100 mM dithiothreitol (DTT) and heated at 95°C for 10 min. Denatured samples were resolved on homemade 15% SDS-PAGE gels for 20 minutes at 80 V then at 140 V for 45-50 min. Gels were stained with PAGE-Blue protein staining solution overnight, destained, and imaged (Biorad ChemiDoc MP).

**Analysis of peptide-Nb crosslinking by liquid chromatography/mass spectrometry (LCMS).** Mass spectrometry data was acquired on a Waters Xevo qTOF LC/MS or an Agilent Affinity II 6130 quadrupole LC/MS instrument. Samples were resolved by reverse-phase LC (Hamilton PRP-h5 column, 5 µM particle size, 300 Å pore size) and analyzed in positive ion mode. For proteins analyzed by mass spectrometry singly charged ions were not observed, so protein intact mass was calculated from analysis of multiply charged ions using the MaxENT algorithm on MassLynx software. Mass spectra corresponding to $Nb_{6E}$ or crosslinked peptide-$Nb_{6E}$ complex were acquired as reference spectra. These reference spectra were used to quantitate the abundances of ions corresponding to $Nb_{6E}$ and crosslinked peptide-$Nb_{6E}$ complex for each time in a kinetic crosslinking reaction. The abundance of $Nb_{6E}$ ions was normalized to the abundance of $Nb_{6E}$ present at $t_0$. The percentage of remaining $Nb_{6E}$ observed for each time point was plotted in GraphPad Prism and fitted to a one-phase decay curve.

**Measurement of crosslinking kinetics using in-gel fluorescence.** Fam-1($NO_2$) peptide and $Nb_{6E}$ crosslinking was conducted in PBS with varying concentrations of Fam-1($NO_2$) and $Nb_{6E}$ in the reaction solution. Conditions tested included 5 µM, 2.5 µM, 1µM, 500 nM, 250 nM of Fam-1($NO_2$) with 100 nM $Nb_{6E}$. At predetermined time points, 20 µL of reaction solution was mixed with unlabeled peptide (6E, final concentration 30 µM) to serve as a competitive inhibitor, effectively quenching the reaction. A positive control consisting of $Nb_{6E}$ labeled with a single fluorescein using sortagging (100 nM, see methods below) was included in each gel. All samples were subsequently denatured by addition of SDS-PAGE loading gel with 100 mM DTT and heated at 95°C for 10 min. All samples were run on 15% SDS-PAGE gels at 80 V for 20 minutes then at 140 V for 45-50 minutes. Samples on the gel were then analyzed for fluorescence via gel imager.

The fluorescence intensity of each band was quantified on ImageJ by outlining a representative rectangular area around a band of interest and measuring the mean brightness values within the selection. This same area was translated to other bands of interest and their mean values also measured. These quantified values were then normalized to the time course sample containing the brightest band.

**Calculation of second-order rate constant of Ac-1($NO_2$) crosslinking.** The second order rate constant for crosslinking was determined as described in precedent literature[4,5]. Band intensities obtained from in-gel fluorescence were plotted as a function of time. These datapoints were then fitted to a one-phase association curve (GraphPad Prism), and the k value associated with each fit was treated as the pseudo-first order rate constant ($k_{app}$). $k_{app}$ values were then plotted against the concentration of crosslinking peptide used in that condition. This resultant plot was fitted to a Michaelis-Menten model and resultant K and $V_{max}$ values determined, where K corresponds to the 1-$Nb_{6E}$ bimolecular $K_D$ value and $V_{max}$ corresponds to the rate constant for labeling ($k_L$). The second order rate constant of crosslinking is calculated as $k_L/K_D$.

**Flow cytometry analysis of fluorescein-labeled crosslinking peptide binding to HEK293 cells.** HEK293 cells stably expressing A2AR($Nb_{6E}$) were cultured as described above. Cells were harvested by trypsinization, which was quenched with the addition DMEM/FBS. Cells were then transferred to a round bottom 96 well plate, pelleted by centrifugation (500 rpm for 3 min.), and resuspended in PBS containing 2% BSA (w/v) (PBS/BSA), and pelleted a second time. The washed cell pellets were then resuspended in PBS/BSA with 100 nM crosslinking Fam-1($NO_2$) or non-crosslinking Fam-1 for 30 minutes on ice. Cells were then washed with PBS/BSA, centrifuged, and resuspended in PBS/BSA containing unlabeled 6E competitor peptide at variable concentrations for 30 minutes on ice. Cells were pelleted then resuspended in PBS/BSA containing Alexafluor647 conjugated anti-fluorescein antibody (1:1000 dilution in PBS/BSA) and incubated for 30 min on ice prior to washing. Washed cells were then resuspended in PBS/BSA for analysis by flow cytometry on a CytoFlex flow cytometer (Beckman Coulter). Live cells were identified based on forward scatter/side scatter profile and staining intensity was monitored in the APC channel. A minimum of 2,000 events corresponding to live cells were recorded.

**Analysis of $Nb_{6E}$ crosslinking on live cells using flow cytometry.** Trypsinized HEK293 stably expressing A2AR($Nb_{6E}$) cells were harvested, transferred into 1.5 microcentrifuge tube, and washed with PBS/BSA. After initial wash, cells were incubated with 100 nM of Fam-1($NO_2$) for indicated durations. At each time point, the crosslinking reaction was quenched with the addition of 100 µL of 10 µM unlabeled 6E competitor. Quenched samples were then diluted to 1 mL with PBS/BSA, pelleted by centrifugation, and media was aspirated. Cells were then resuspended stained with AF647-labeled secondary antibody (Jackson ImmunoResearch, 200-602-037) and analyzed as above. Flow cytometry histograms were used to calculate median fluorescence intensity (MFI) values for the APC channel in each sample. MFI values were normalized to internal controls and averaged among replicate samples.

**Fluorescence microscopy analysis of nanobody staining.** A 4-chamber glass slide (Lab-Tek II Chamber SlideTM) was treated with 10% polylysine lysine solution (500 µL/chamber) for 5 minutes, rinsed with molecular-biology grade water and allowed to dry for 1 hour. Trypsinized HEK293 cells were harvested were transferred to a 4-well glass slide and left to incubate in DMEM/FBS at 37°C overnight at which point a confluency of approximately 90% was reached. The medium was aspirated, cells were washed once with PBS/BSA (1 mL) and placed on ice. Cells were stained with 500 µL of indicated solutions for 30 minutes. After incubation, chambers were subsequently washed with PBS/BSA and then fixed with 4% paraformaldehyde solution for 15 minutes. The chambers were then washed with PBS/BSA. Following the wash, the chamber walls were removed and mounting solution containing DAPI was applied to each chamber (ProLong Glass Antifade Mountant with NucBlue). Chambers were enclosed with a coverslip (Gold Seal Cover Glass 24x50 mm No. 1 1/2) and allowed to cure overnight in the dark.

Stained cells were imaged using a Nikon Eclipse 50i microscope coupled to a Cool Snap ES2 CCD camera (Photometrix). Images were acquired at 20x magnification using various filter channels (DAPI, Fluorescein, and Rhodamine). For visualization of each stain, the following exposure times were used: DAPI (5 sec.), Fluorescein (30 sec.), and Rhodamine (30 sec.) Images were processed with ImageJ (FIJI package).  and Fluorescein signal brightness normalized for each sample (minimum: 20, max: 100). Images acquired at each channel were merged to obtain composites.

**Protein labeling via sortagging.** Sortagging reactions were comprised of the following components: protein bearing a sortase recognition motif (LPETGG) followed by a hexa histidine tag at the C-terminus (20-200 µM final concentration), triglycine-probe conjugates (500-1000 µM final concentration), and Sortase 5M (10-20 µM final concentration). Reactions were performed in sortase buffer (10 mM $CaCl_2$, 50 mM Tris, 150 mM NaCl, pH 7.5) and shaken at 12°C overnight. After incubation, the reaction was incubated with nickel NTA beads to capture Sortase 5M and unreacted starting protein. Uncaptured material was further purified using disposable desalting columns to remove triglycine conjugates (Cytiva PD-10 Sephadex™ G-25M). Eluents were monitored for absorbance at the fluorescent probe absorbance wavelength (tetramethylrhodamine: 555 nm, Fluorescein: 494 nm), 220 nm, and 280 nm for the presence of protein conjugate. Fractions containing product were combined then concentrated by spin filtration (Amicon Ultra 0.5 mL Centrifugal Filters 10,000 NMWL).

**Western Blotting.** Cultured HEK293 cells were trypsinized using standard protocols described above and pelleted by centrifugation. The media was then aspirated, the cells resuspended in PBS, and transferred into 1.5 mL microcentrifuge tubes on ice where labeling was performed. Cells were then labeled with desired crosslinking peptides. Post-labeling, cells were washed with PBS, pelleted, then resuspended in 40 µL of 1% NP40 in PBS for 20 min to lyse cells. Samples were then centrifuged for 10 min (14,000 RPM, 4°C) to pellet nucleus and other insoluble cellular debris. The supernatant was transferred to a new 1.5 mL centrifuge tube containing SDS-PAGE sample buffer with 100 mM DTT and incubated at 60°C for 10 minutes. Samples were run on SDS-PAGE gels (homemade 15% SDS-PAGE gels or 4-20% Mini-Protean TGX gels, Biorad #4561094) at 140 V then subsequently transferred to a PDVF membrane by transfer blotting (1.3 A, 25V, 7 min).

Post-transfer, the PDVF membrane was blocked overnight at 4°C with non-fat milk solution (PBS, 0.1% Tween 20, 5% w/v dry milk). After overnight blocking the PDVF membrane was washed with 0.05% Tween20 in PBS (PBST). The membrane was then stained with Peroxidase IgG Fraction Monoclonal Mouse Anti-Fluorescein in PBS (Jackson ImmunoResearch 200-032-037, 1/3000 dilution) for 1 hour, then washed 3x with PBST. Antibody staining was then visualized using Pierce ECL Western blotting substrate on a gel imager.

GAPDH detection was performed by inactivating membrane-bound HRP by treating with 0.1 mM sodium azide in 2%BSA/PBS overnight at 4°C. The membrane was then washed with PBST (3x) and then treated with 10 mL of Anti-GAPDH antibody conjugated with HRP (BioLegend #607903, 1:3000 dilution in PBS) and incubated for 45-minutes. Excess Anti-GAPDH antibody washed away with PBST and the membrane was imaged as above.

**Surface plasmon resonance (SPR) binding experiments.** SPR measurements were performed on a GE Biacore T100 – T200 Sensitivity Enhanced Instrument using a Cytiva Series S Sensor Chip SA (immobilized streptavidin). Ligand ($Nb_{6E}$-biotin) was prepared using standard sortase ligation protocols described above and diluted to a concentration of 80 µg/mL. The SA chip was conditioned using successive treatments with 1) 1M NaCl, 50 mM NaOH 2) 50% isopropanol, 50 mM NaOH, 1M NaCl.

Analyte samples (6E and analogues) were prepared via two-fold serial dilutions in PBST ranging from 1.6 µM to 3.125 nM. Analyte was flowed over the chip at 50 µL/min with a contact time 60 seconds, and a dissociation time of 120 sec. Regeneration step (to dissociate 6E peptides) included of consecutive washes of 10 mM glycine solution (pH 1.5). Sensorgrams were fitted using the Biacore T200 evaluation software to a 2-state model, with local Rmax. Raw sensorgrams and their respective fits were exported as ASCII files and regraphed on GraphPad prism.

**Mass spectrometry-proteomic analysis of Ac-1-$Nb_{6E}$ crosslinking.** $Nb_{6E}$ was labeled using Ac-1($NO_2$) at a 4-fold molar excess. Crosslinked product was purified from excess crosslinking peptide using a PD10 column into PBS for subsequent analysis by mass spectrometry. This sample was diluted in buffer containing 50 mM Tris, 100 mM hydroxyproline, 8M urea, and 5 mM DTT and heated to 40° C for 5 minutes and incubated at room temperature for 15 minutes. Chloroacetamide was then added to the sample (15 mM) and the sample was incubated for 30 minutes in the dark at room temperature. Excess chloroacetamide was then scavenged by addition of 10 mM betamercaptoethanol incubation for an additional 15 minutes at room temperature. For the Lys-C digest 20 µL of 1M Tris and 80 µL of water was then added (dilution factor of two). For the Trypsin digest 40 µL of 1M Tris and 160 µL of water was then added (dilution factor of four). 0.4 µg Trypsin or Lys-C was added to initiate digestion (20:1 subject:protease ratio). After 2 hours of digestion the sample was acidified by adding formic acid to 1.6%, chilled in an ice water bath, and then applied to a stack of in house Empore C8/C18 stage tips[6], which were loaded in a swinging bucket rotor at 300 G at 5° C. The tip stack was washed with 3 x 150 µL 1.6% formic acid, 50 mM ammonium acetate. Sample was then eluted with 0.4% formic acid 40% acetonitrile solution and then 1.6% formic acid 80% acetonitrile solution through centrifugation at 300 G. Samples were dried under nitrogen with heating and re-suspended in 0.1% formic acid 2% acetonitrile.

LC/MS/MS analysis was carried out using a Thermo Lumos mass spectrometer operating in a high/high approach together with a Thermo nLC1000 with aftermarket modification to address known rotor issues (VICI). Experiments were carried out using a 500 mm Easy Spray (ES903) column at a temperature of 50° C. Data was collected as a single MS1 spectrum (resolution setting 120K) followed by up to a 3 second window collecting MS2 spectra (15K, HCD 30) with targets selected based on charge state (3-5) The nLC-1000 was operated at 150 nL/minute with A solvent 0.1% formic acid in water and B solvent 0.1% formic acid in 93.75% acetonitrile in water. The gradient run was 2-35% in 80 minutes, 35-80% in 20 minutes, and holding at 80% for 20 minutes.

MaxQuant (1.6.10.43)[7] was used to process the data searching the contaminants database, a file which contained the sequence of the protein construct, and a mock sequence which would produce the crosslinked peptide fragments. Modifications (all variable) included cysteine carbamidomethylation, methionine oxidation, modification with the described peptide and cross linker (with various targetable amino acids).

**Time Course Hydrolysis of Ac-1(NO2)** Purified 1 mM stock of Ac-1($NO_2$) in DMSO was dissolved in PBS to a final concentration of 50 µM and incubated at room temperature. At predetermined time points, 20 uL of reaction solution was diluted with 80 uL of DI $H_2O$. 90 uL of this solution was run on an analytical HPLC using a Phenomenex Aeris Peptide XB-C18 Analytical column (particle size 5 µM, 100 Å pore size) with a linear gradient of 10-80% acetonitrile in water 0.1% trifluoroacetic acid. Peaks corresponding with the starting material were integrated and normalized with respect to the initial timepoints, to obtain % of starting material remaining.

**Synthetic protocols:**

**Maleimido-phenol ester synthesis.** 6-maleimidohexanoic acid (Alfa Aesar, 46384) was dissolved in DMF and mixed with 1 equivalent diisopropylcarbodiimide (DIC). Phenol derivatives (5 equivalents) were dissolved in a separate vial in DMF with diisopropylethylamine (DIPEA, 10 equivalents) which was then added drop wise to the activated 6-maleimidohexanoic acid. The reaction mixture was shaken overnight at 4°C. Precipitate was removed by centrifugation and product was purified by reverse-phase HPLC (C8 column, gradient 20-90% acetonitrile in water with 0.1% trifluoroacetic acid). Product stock solutions were made at a concentration of 100 mM in DMSO.

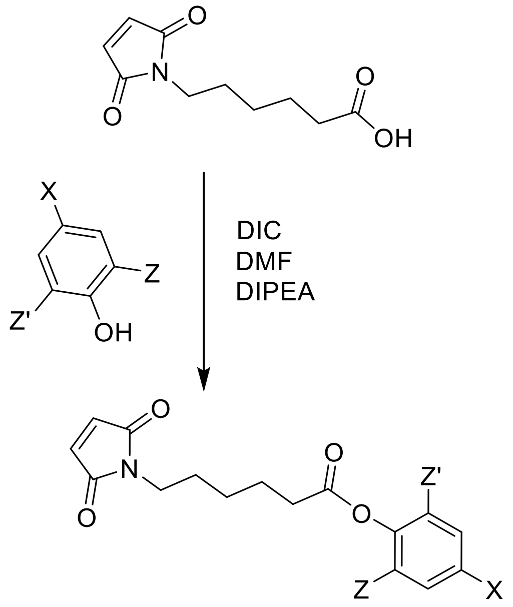


X = $NO_2$; Z/Z' = H (**Mal-$NO_2$**)
X =H; Z/Z' = Br (**Mal-DiBr**)

**Activated 6E phenolic esters via Cys-maleimide chemistry.** Purified peptides were dissolved in DMSO (10 mM) and mixed with 3-5 equivalents of maleimide-phenol ester. Concentrated phosphate buffer (1 M, pH 7.5) was then added (final concentration 100 mM) to adjust reaction pH. The reaction was shaken at room temperature for 1 h and the product was purified by reverse-phase HPLC (C18 column, gradient 20-90% acetonitrile in water with 0.1% trifluoroacetic acid), lyophilized and dissolved in DMSO (1 mM stock) prior to use.

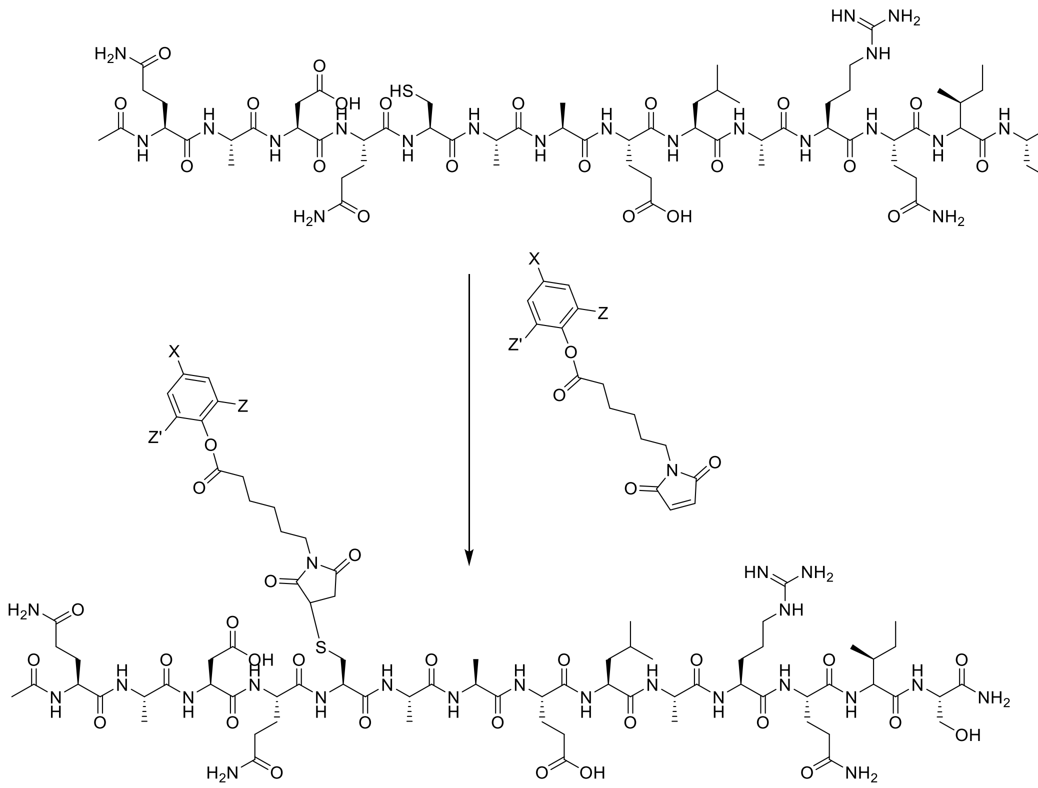

X = $NO_2$; Z/Z' = H (**1-$NO_2$**)
X =H; Z/Z' = Br (**1-DiBr**)

**CGS-alkyne**. CGS21680 (Cayman Chemical, # 17126) was dissolved in dimethylformamide (DMF) and diisopropylcarbodiimide (1 equivalent) was added. To this mixture 2 molar equivalents of $NH_2$-$PEG_4$-alkyne was dissolved in DMF and was added (Click Chemistry Tools, # TA101-100). Next, diisopropylethylamine (5 molar equivalents) was added. The reaction was shaken at 10°C overnight. The reaction was purified by reverse-phase preparatory HPLC (C18 column, gradient 20-90% acetonitrile in water with 0.1% trifluoroacetic acid).

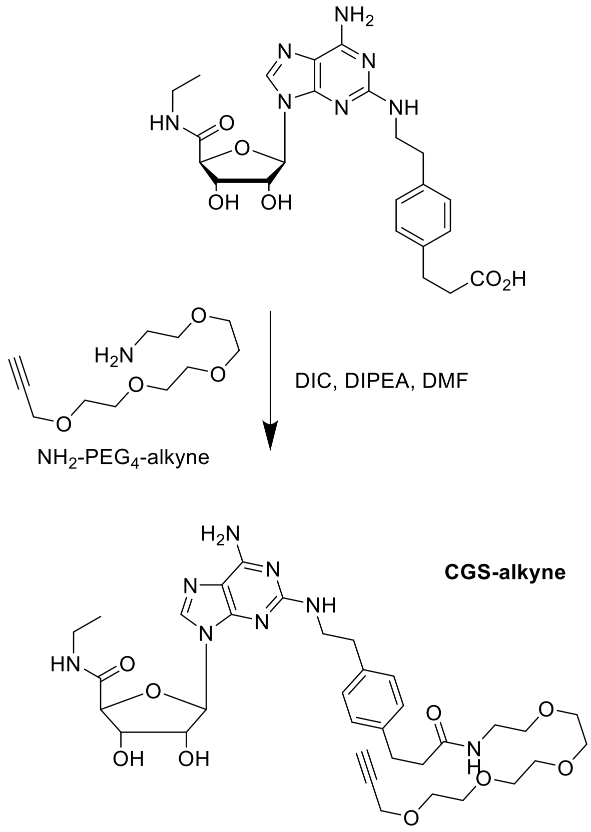


**Copper-catalyzed click chemistry (CGS-6E-A5 and CGS-1)**

Azide-alkyne conjugation was performed as previously described[8] with some modifications. Azide-6E peptides were dissolved in DMF (5 mM) and CGS-alkyne was added at a 3-fold excess (15 mM). $CuSO_4$ heptahydrate was dissolved in water and mixed with THPTA (Click Chemistry Tools, #1010-100) at a 1:5 molar ratio to prepare a 10x stock solutions (1 mM $CuSO_4$, 5 mM THPTA). The pre-mixed copper-THPTA solution was added then added to the DMF solution of azide and alkyne. A fresh stock solution of 100 mM sodium ascorbate was prepared in water and diluted 1:20 into the reaction solution to initiate the reaction (final concentration of 5 mM). The reaction was monitored by LC/MS and additional aliquots of premixed $CuSO_4$/THPTA and sodium ascorbate were added over the course of 12-48 h to promote conversion to product. The reaction was purified by reverse-phase HPLC, lyophilized, and dissolved in DMSO at a concentration of 1 mM for use in biological assays.

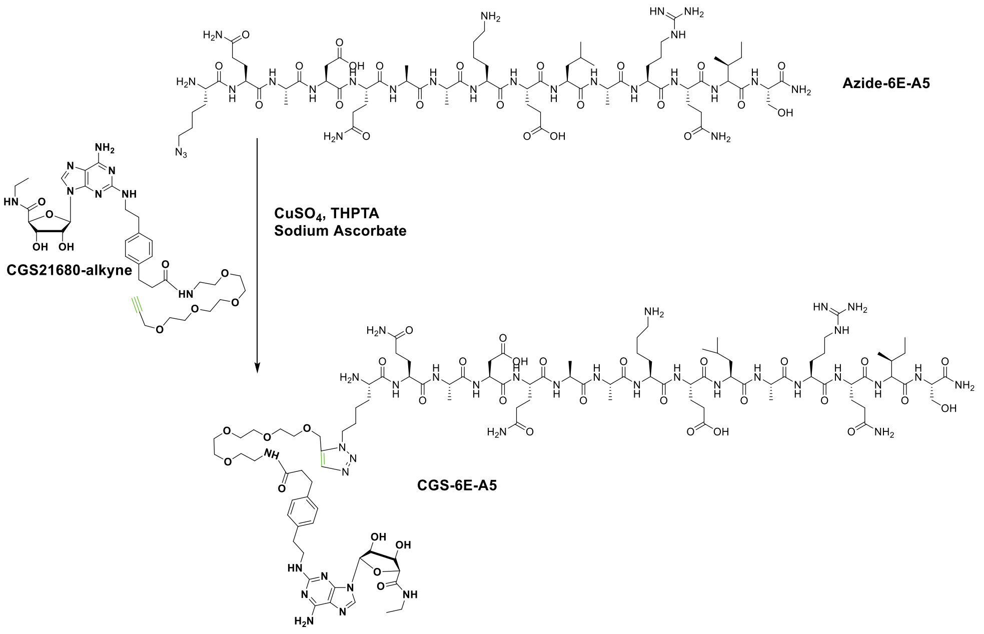


### CGS-6E-A5

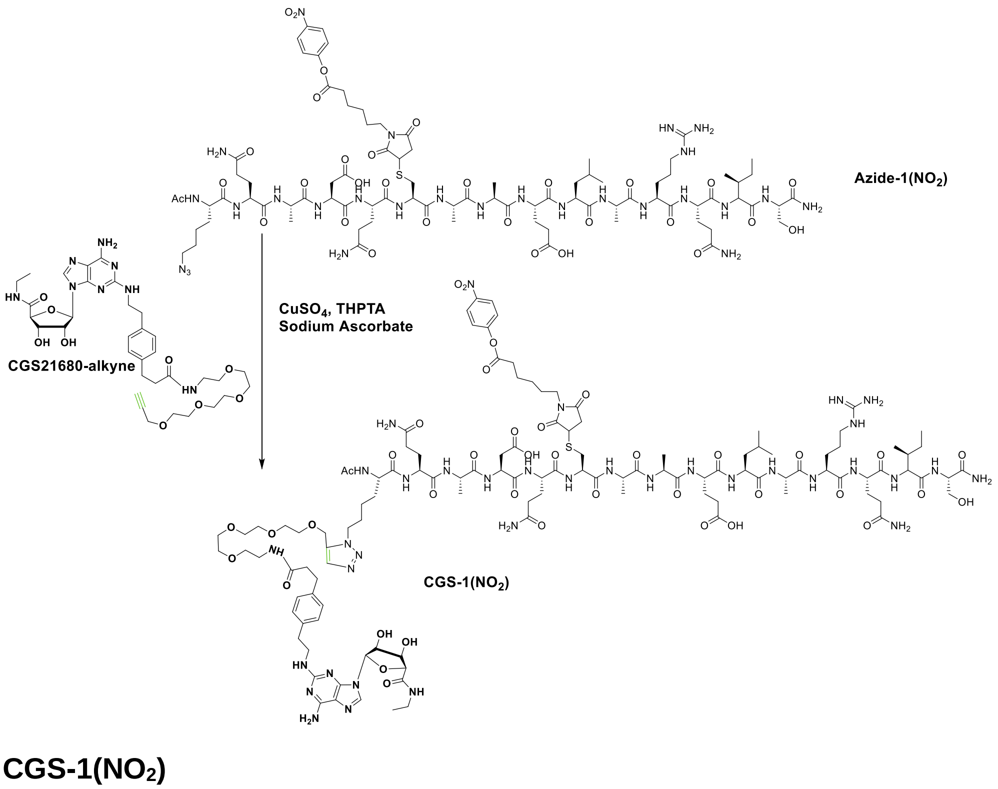


### CGS-1($NO_2$)

**References.**
